# Mining the biosynthetic landscape of lactic acid bacteria unearths a new family of RiPPs assembled by a novel type of ThiF-like adenylyltransferases

**DOI:** 10.1101/2024.05.13.593846

**Authors:** Mengjiao Wang, Mengyue Wu, Meng Han, Xiaogang Niu, Aili Fan, Shaozhou Zhu, Yigang Tong

## Abstract

Ribosomally synthesized and post-translationally modified peptides (RiPPs) are chemically diverse natural products of ribosomal origin. These peptides, which frequently act as signals or antimicrobials, are biosynthesized by conserved enzymatic machineries, making genome mining a powerful strategy for unearthing previously uncharacterized members of their class. Herein, we investigate the untapped biosynthetic potential of Lactobacillales (i.e. lactic acid bacteria) – an order of Gram-positive bacteria closely associated with human life, including pathogenic species and industrially-relevent fermenters of dairy products. Through genome mining methods, we systematically explored the distribution and diversity of ThiF-like adenylyltransferases-utilizing RiPP systems in lactic acid bacteria and identified a number of unprecedented biosynthetic gene clusters. In one of these clusters, we found a previously undescribed group of macrocyclic imide biosynthetic pathways containing multiple transporters and may be involved in potential quorum sensing (QS) system. Through in vitro assays, we determined that one such adenylyltranferase specifically catalyzes the intracyclization of its precursor peptide through macrocyclic imide formation. Incubating the enzyme with various primary amines revealed it could effectively amidate the C-terminus of the precursor peptide. This new transformation adds to the growing list of Nature’s peptide macrocyclization strategies and expands the impressive catalytic repertoire of the adenylyltransferase family. The diverse RiPP systems identified herein represent a vast, unexploited landscape for discovery of novel class of natural products and potential QS systems.

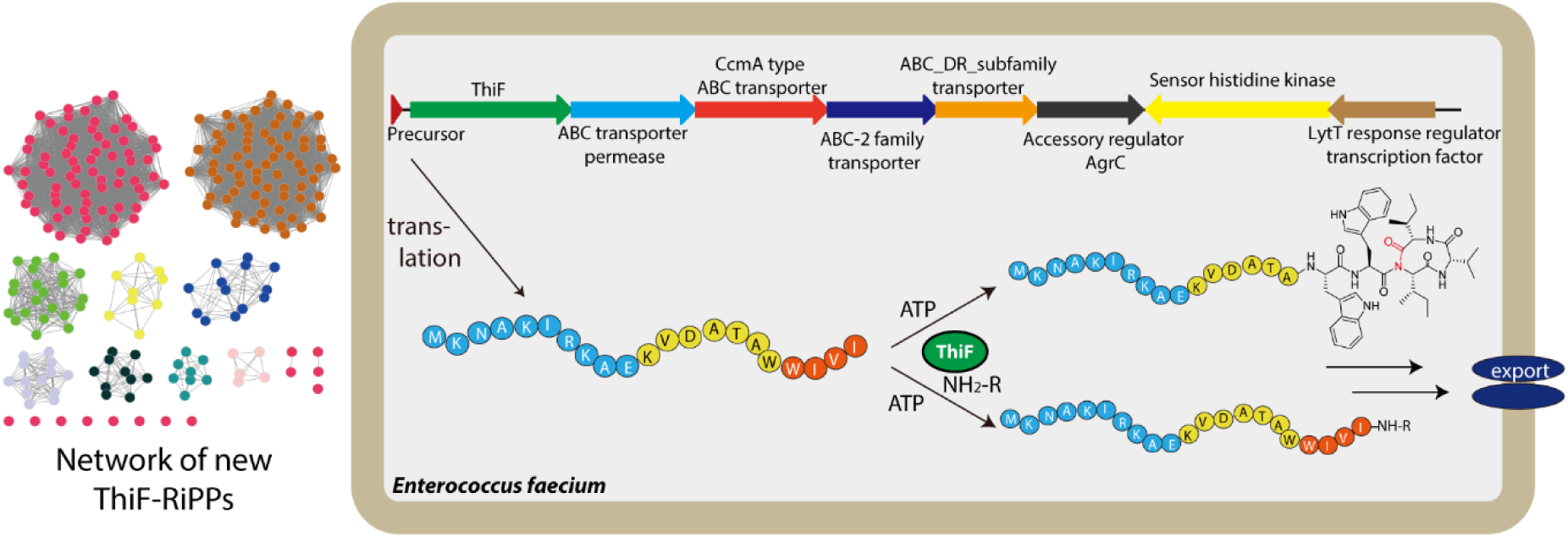

(For Table of Contents use only)

## INTRODUCTION

Ribosomally synthesized and post-translationally modified peptides (RiPPs) are a growing family of natural products that exhibit remarkable chemical diversity in spite of their ribosomal origin. On account of their complex structures, these peptides carry out an array of medically relevant functions, serving as antibiotics, anticancer agents, cofactors, signals, and toxins.*^1, 2^*

Typically, RiPPs are biosynthesized as precursor peptides consisting of two parts: an N-terminal leader sequence, which is bound and recognized by processing enzymes, and a C-terminal core sequence, which is matured by that machinery.*^3^* Often, the leader sequence is proteolytically cleaved during processing, freeing the core scaffold for further embellishment by option tailoring enzymes prior to transporter-mediated export from the cell. Much of the chemical (and thus functional) diversity of RiPPs hinges on cyclization reactions that substantially alter the three-dimensional shape of the peptide, resulting in e.g. proteolytic stability and/or affinity for specific molecular targets (Fig. 1).*^4^*

**Figure 1.**
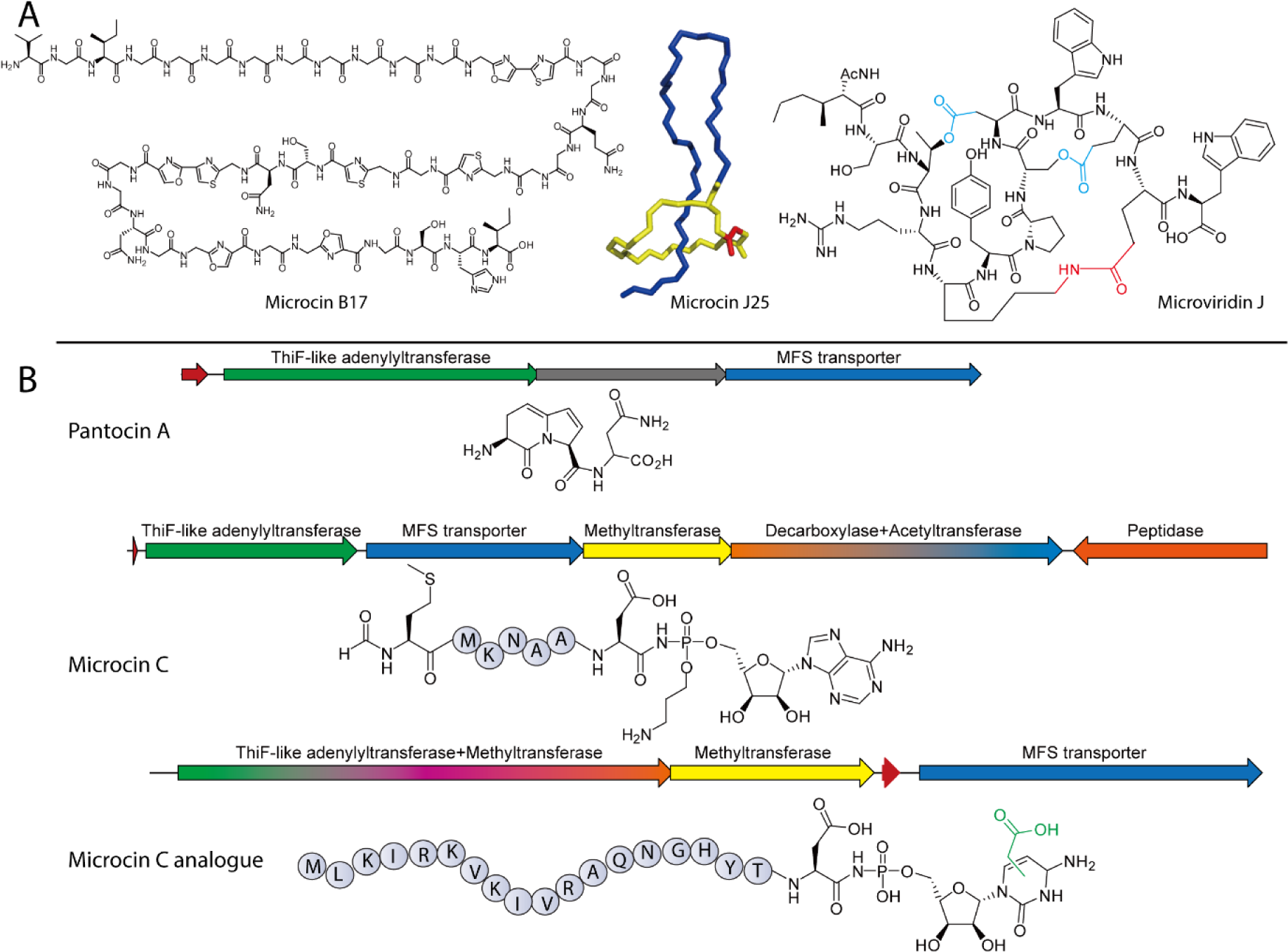
(A) Representative structures of known cyclic peptides with isopeptide bonds, (B) Known RiPPs biosynthetic gene clusers containing ThiF enzymes including pantocinA, microcin C and microcin C analogue.

ATP/Mg^2+^-dependent activation of the precursor peptide with phosphate or AMP is a common strategy for forming cyclic moieties during RiPP biosynthesis.*^4^* For example, it is employed to generate the heterocycles of thiazole/oxazole-modified microcins (LAPs; e.g. microcin B17),*^5, 6^* the cyclic thioethers of lanthipeptides (e.g. lacticin 481),*^7^* and the macrolactams of microviridins and lasso peptides (Fig. 1A).*^8–10^*

A distinctive set of enzymatic machineries has evolved to install the characteristic cyclic features in each of these RiPP subfamilies. For instance, the cyclodehydration reactions in LAP biosynthesis require a protein homologous to the adenylyltransferases ThiF, MoeB, and ubiquitin-activating enzyme E1 that bind and adenylate their substrates during thiamine or molybdopterin biosynthesis and protein ubiquitination, respectively.*^11–13^* Unlike ThiF, MoeB, and E1, however, the ATP-binding site is perturbed and no ATP consumption has been demonstrated for the homologs involved in LAP assembly(e.g., McbB for microcin B17); instead, the latter appear to bind the peptide substrate, assisted by an N-terminal RiPP precursor peptide recognition element (RRE),*^14^* and deliver it to an ATP-dependent YcaO-like enzyme for cyclization (e.g. McbD for microcin B17).*^15–18^*

Conversely, adenylyltransferases from other RiPP systems like those of pantocin A and microcin C function more like ThiF, MoeB, and E1 by directly activating the C-termini of their substrates with AMP, again with support from RREs (we refer to these homologs hereafter as ThiF-like adenylyltransferases, or TLATs).*^11, 19^* Past research correlated the RiPP gene cluster *paaPABC* from *Pantoea agglomerans* with the biosynthesis of pantocin A, a bicyclic tripeptide (EEN) that exerts its antibiotic activity through inhibition of l-histidinol phosphate aminotransferase (Fig. 1B).*^20, 21^* More recent *in vitro* investigations revealed that TLAT PaaA promotes ATP-dependent intramolecular cyclization through two dehydrations and one decarboxylation of the Glu pair in the middle of precursor peptide PaaP, leading to formation of the bicyclic core of pantocin A.*^19^* α-Ketoglutarate-dependent iron oxidase PaaB, an unidentified peptidase, and ATP-binding cassette (ABC) transporter PaaC then catalyze dehydrogenation, proteolysis, and efflux, respectively.*^19, 20^*

Microcin C, on the other hand, is an aspartyl-tRNA synthetase-targeting antibiotic produced by a plasmid-born *mccABCDEF* gene cluster in *Escherichia coli* (Fig. 1B).*^22, 23^* This *N*-formylated heptapeptide (MRTGNAD) is linked at its C-terminus to AMP via an N-P bond, installed by TLAT MccB in two ATP-dependent reactions: intramolecular cyclization between Asn and the C-terminus to form a peptidyl-succinimide intermediate, followed by adenylation of the succinimidyl N atom and hydrolytic opening of the succinimide ring.*^24^* Finally, *S*-adenosylmethionine (SAM)-dependent methyltransferase and pyridoxyl 5ꞌ-phosphate (PLP)-dependent decarboxylase homologs MccD and MccE attach propylamine to the phosphate group, major facilitator superfamily (MFS) transporter MccC exports the compound from the cell, and serine protease MccF provides immunity.*^22, 23, 25–27^*

Recently, a microcin C analogue consisting of a 19-residue peptidyl region and a carboxymethyl-modified cytidine moiety in place of the C-terminal adenosine was discovered from *Bacillus amyloliquefaciens* (Fig. 1B).*^28^* The MccB homolog from this pathway, was shown to accept all four NTPs *in vitro*, albeit with striking preference for CTP, in agreement with the peptidyl-cytidylate isolated from bacilli. Furthermore, a C-terminal extension to MccB, identified as a SAM-dependent methyltransferase homolog, was confirmed to install the carboxymethyl group in conjunction with carboxy-SAM synthetase homolog MccS. As in the *E. coli* gene cluster, a gene coding for an MFS transporter likely responsible for export of the antibiotic was observed.

Severinov and co-workers previously identified and/or functionally validated additional homologs of PaaA, MccB, and associated precursor peptides encoded not only Gram-negative bacteria but bacillota and cyanobacteria.*^29, 30^* This finding suggests that many more TLATs, and thus pantocin A-and microcin C-like RiPPs, remain to be discovered. More recently, Seyedsayamdost *et al*. have shown that ThiF-like adenylyltransferase/cyclase is also involved in the biosynthesis of RiPPs tailored by radical S-adenosylmethionine (RaS) enzymes in *Streptococci*. This enzyme can generate a C-terminal Glu-to-Cys thiolactone macrocycle, further demonstrating the versatility of these enzymes.*^31^*

Given the few but diverse TLAT-RiPP systems that have been characterized, we decided to seek out further examples. We began by mining for homologs of MccB in the genomes of lactic acid bacteria. On account of their relatively compact genomes, this order of Gram-positive bacteria, consisting of human commensals and opportunistic pathogens as well as food industry mainstays,*^32–34^* has been underexploited for natural product discovery.*^35, 36^* Our efforts uncovered an extensive collection of BGCs that, based on their unique architectures, appear to encode RiPPs decorated with previously undescribed post-translational modifications. A second stage of mining performed without limiting genomes to lactic acid bacteria revealed that RiPP-associated TLATs are employed liberally across the bacterial kingdom. We selected one TLAT identified from *Enterococcus faecalis* FDAARGOS_397, EnfB, for further characterization, which led to the discovery of enterofaecin, a microcin-like peptide containing a macrocyclic imide distinct from succinimide. We also found that, in the presence of EnfB, enterofaecin undergoes C-terminal amidation by various primary amines. The association of diverse ATP-binding cassette (ABC) transporter and a novel QS system indicated that this biosynthetic logic might play important physiological roles such as virulence or interspecies communication. Our studies open an adit to a vast network of RiPPs with unprecedented catalytic inventory and yet-to-be explored biological functions.

## Materials and Methods

### Bioinformatics studies

Genome mining studies were performed using the web-based bioinformatic tool PSI-BLAST, with MccB^Eco^ (AAY68495.1) as the query sequence.*^37^* Algorithm parameters were set as follows: maximum target sequences, 1000; expect threshold, 10; matrix, BLOSUM62. The organism was limited to either Streptococcus (taxid: 1301) or Enterococcus (taxid: 1350). This resulted in 2000 TLAT-encoding loci.

Sequence similarity network (SSN) analysis was carried out using the Enzyme Function Initiative-Enzyme Similarity Tool (EFI-EST) with input fasta files containing all protein sequences to be analyzed.*^38^* After the initial datasets were generated, we used an alignment score that would correspond to 20% sequence identity for outputting and interpretation of the SSN. The resulting xgmml file was imported into Cytoscape 3.8.2.*^39^* Each node in the network corresponds to a single protein sequence and each edge represents a pairwise connection between two proteins based on the specified sequence identity cutoff. The networks were visualized using the yFiles organic layout option of Cytoscape 3.8.2.

Biosynthetic gene clusteres were annotated with consensus sequences generated by Geneious 10.2.2.*^40^*

### Gene synthesis and materials

The gene encoding EnfB (WP_098041451) from *Enterococcus faecium* strain FDAARGOS_397 was synthesized by Synbio Technologies (Suzhou). NTPs were purchased from Sigma Aldrich. Peptides were synthesized by Chinapeptides. Q5 High-Fidelity DNA Polymerase was obtained from New England Biolabs. DNA manipulation and purification kits were purchsed from Tiangen (China). Kanamycin, chloramphenicol, and other chemicals were obtained from commercial sources. Primers listed in Table S2 were purchased from RuiBiotech® (Beijing, China).

### Cloning, expression, and purificaiton of EnfB and its variants

Through Gibson assembly (ThermoFisher), the gene encoding EnfB (synthesized by TransGen Biotech, China) was ligated into pET30a (Invitrogen, USA) at the *Nde*I and *Xho*I restriction sites, with a His_6_ tag introduced at the C-terminus. The resulting mixture was introduced into chemically competent TOP10 *E. coli* cells by heat shock, and cells were subsequently inoculated on solid lysogeny broth (LB) medium containing kanamycin (50 μg/mL). The plasmid was isolated from a single transformant in LB medium with kanamycin (50 μg/mL) using DNA purification kits purchased from Tiangen (China) and verified by DNA sequencing(RuiBiotech, China).

The correct plasmid was introduced into chemically competent Transetta(DE3) *E. coli* cells ((Weidi, China)) by heat shock and plated on solid LB medium supplemented with kanamycin (50 µg/mL) and chloramphenicol (34 µg/mL). LB medium (50 mL) containing kanamycin (50 µg/mL) and chloramphenicol (34 µg/mL) was inoculated with a single colony for 16 h as a preculture. Next, 2 mL of the preculture was transferred to 200 mL LB medium (with 50 µg/mL kanamycin and 34 µg/mL chloramphenicol) and incubated at 37 °C and 200 rpm to an optical density at 600 nm of 0.6-0.8. Gene expression was induced with 0.1 mM IPTG at 16 °C and 150 rpm for 20 h. After harvesting cells by centrifugation (7000 rcf, 15 min, 4 °C), the resulting pellets were resuspended in 50 mL Lysis Buffer (20 mM Tris, 500 mM NaCl, 30 mM imidazole, 10% glycerol, pH 7.4) containing hen egg white lysozyme (1 mg/mL) and incubated on ice for 10 min. Cells were lysed by an ultra high pressure crusher(JNBIO, China), and insoluble components were removed by centrifugation (7000 rcf, 1 h, 4 °C). The supernatant containing recombinant protein was then filtered and loaded onto a Ni-NTA gravity flow column at 4 °C. After removing nonspecifically bound proteins with Wash Buffer (20 mM Tris, 500 mM NaCl, 50 mM imidazole, 10% glycerol, pH 7.4), recombinant EnfB was released from the column with Elution Buffer (20 mM Tris, 500 mM NaCl, 500 mM imidazole, 10% glycerol, pH 7.4). Purified protein was exchanged into Buffer S (10 mM Tris, 200 mM NaCl, 5 mM DTT, 10% glycerol, pH 7.8) and concentrated to 5 mL in an Amicon Ultra Centrifugation Filter (10 kDa), then further purified on a Superdex 200 column (GE) in Buffer S with a FPLC system (Bio-Rad). After size-exclusion chromatography, EnfB-containing fractions were selected by analysis on a 12% (w/v) SDS-PAGE gel, concentrated in an Amicon Ultra Centrifugation Filter, flash-freezed in N_2_ and stored at −80 °C.

### *In vitro* assays with EnfB

Chemically synthesized EnfA (200 μM, Chinapeptides) was dissolved either in phosphate buffer (0.2 M phosphate buffer, pH 7.0, 5 mM NTP, 10 mM MgCl_2_) or Tris buffer (0.1 M Tris buffer, pH 7.0, 5 mM NTP, 10 mM MgCl_2_) and incubated with EnfB (1.75 μM) in 100 μL reactions at 32 °C for 3 h. Reactions were quenched through addition of 1 μL TFA and cleared by centrifugation (12,000 rcf, 5 min). Supernatants (10 μL) were then subjected to HPLC (controller, Shimadzu CMB-20A; diode array detector, Shimadzu SPD-20A; pumps, Shimadzu LC-20A) on a reverse-phase column (SilGreen, particle size 5 μm, 12 nm, 4.6 × 250 mm) at a flow rate of 0.8 mL/min. The following gradient of solvent A (water containing 0.1% TFA) and solvent B (acetonitrile containing 0.1% TFA) was used: 20% B for 2 min, up to 45% B in 8 min, up to 95% B in 10 min, 95% B for 5 min, and down to 20% B in 7 min. Substrate and product concentrations were ascertained from UV peak area counts (214nm).

For time-dependent conversion of EnfA by EnfB, *in vitro* assays were carried out as above (with ATP as the NTP) for varying durations (0, 1, 2, 3, 4, 6, or 10 h) before quenching, clearing, and analysis. Similarly, the effect of temperature on EnfB activity was investigated over a range of 10 to 60 °C (10, 20, 30, 40, 45, 50, 55, or 60 °C) for 3 h at 600 rpm. The effect of pH on EnfB activity was likewise probed in Reaction Buffer at pH values between 4.0 and 10.0 (adjusted through addition of NaOH or HCl) for 3 h at 50 °C. Finally, the kinetic constants of EnfB were measured similarly by adding EnfB (5.46 μM) to 100 μL Reaction Buffer containing varying concentrations of EnfA (10, 20, 30, 40, 60, 100, 200, or 500 μM) for 3 h at 50 °C.

EnfA variants were synthesized (Chinapeptides) and, along with various primary amines, used to probe the substrate specificity of EnfB. The standard *in vitro* assays were performed as above, except with 2.72 μM EnfB and 200 μM EnfA variant (see Fig. 5A) or one of amines **1**-**6** (see Fig. 5B) in place of EnfA for 12 h at 32 °C and 600 rpm before quenching, clearing, and analysis.

**Figure 2.**
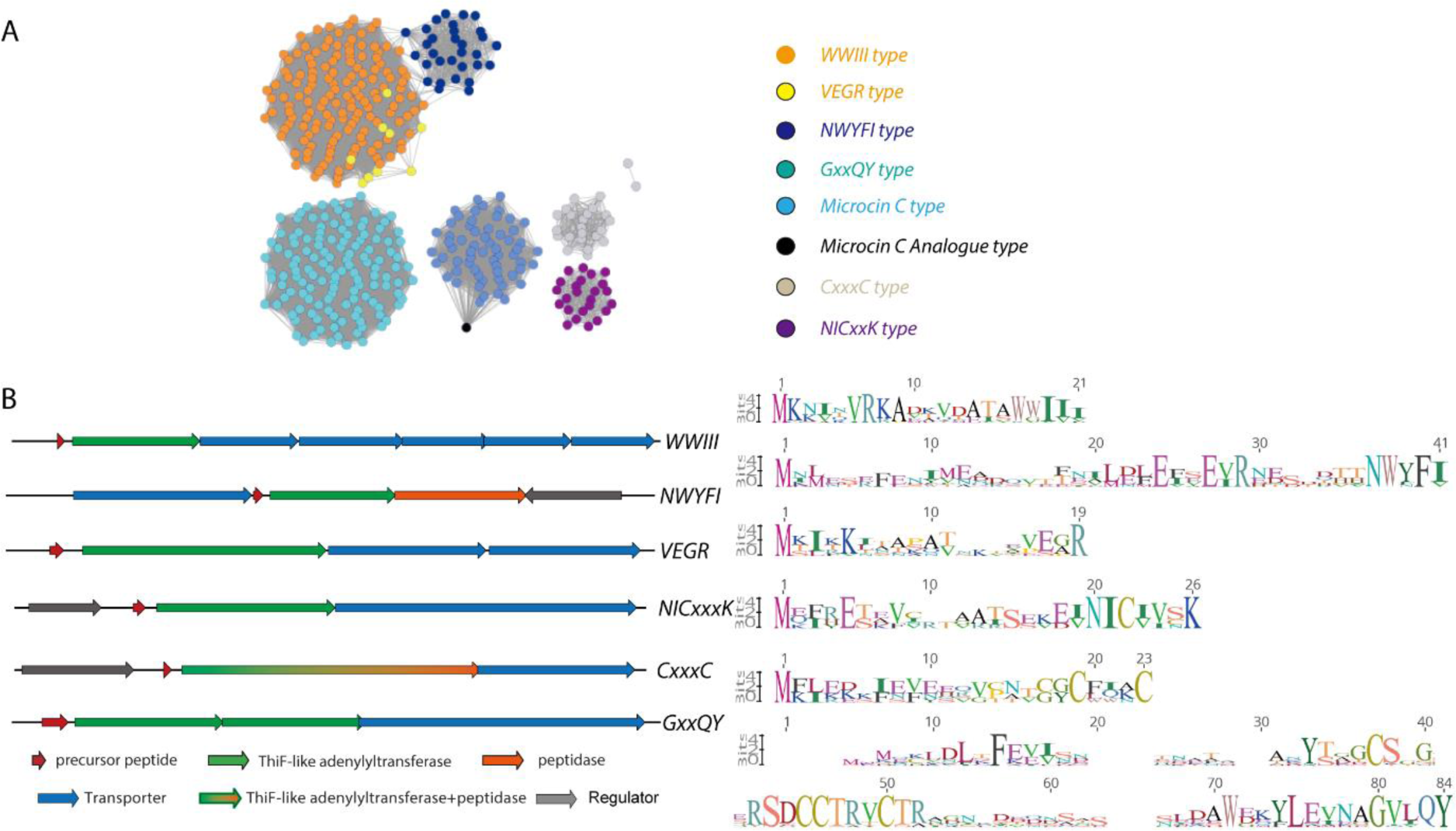
ThiF-RiPP Network in Lactobacillales. (A) sequence similarity network of ThiF-RiPP gene clusters from 8 groups (edge % identity of 20) based on the sequence of the ThiF genes. Color-coding indicates the distribution of each subfamily among Lactobacillales strains. The subfamilies are named based on conserved motifs within each precursor peptide. (B) Representative biosynthetic gene cluster for each of the sub-families in panel A. The genes are color-coded as indicated on the figure. Precursor peptide logo plots for ThiF-RiPP subfamilies are shown on the right.

**Figure 3.**
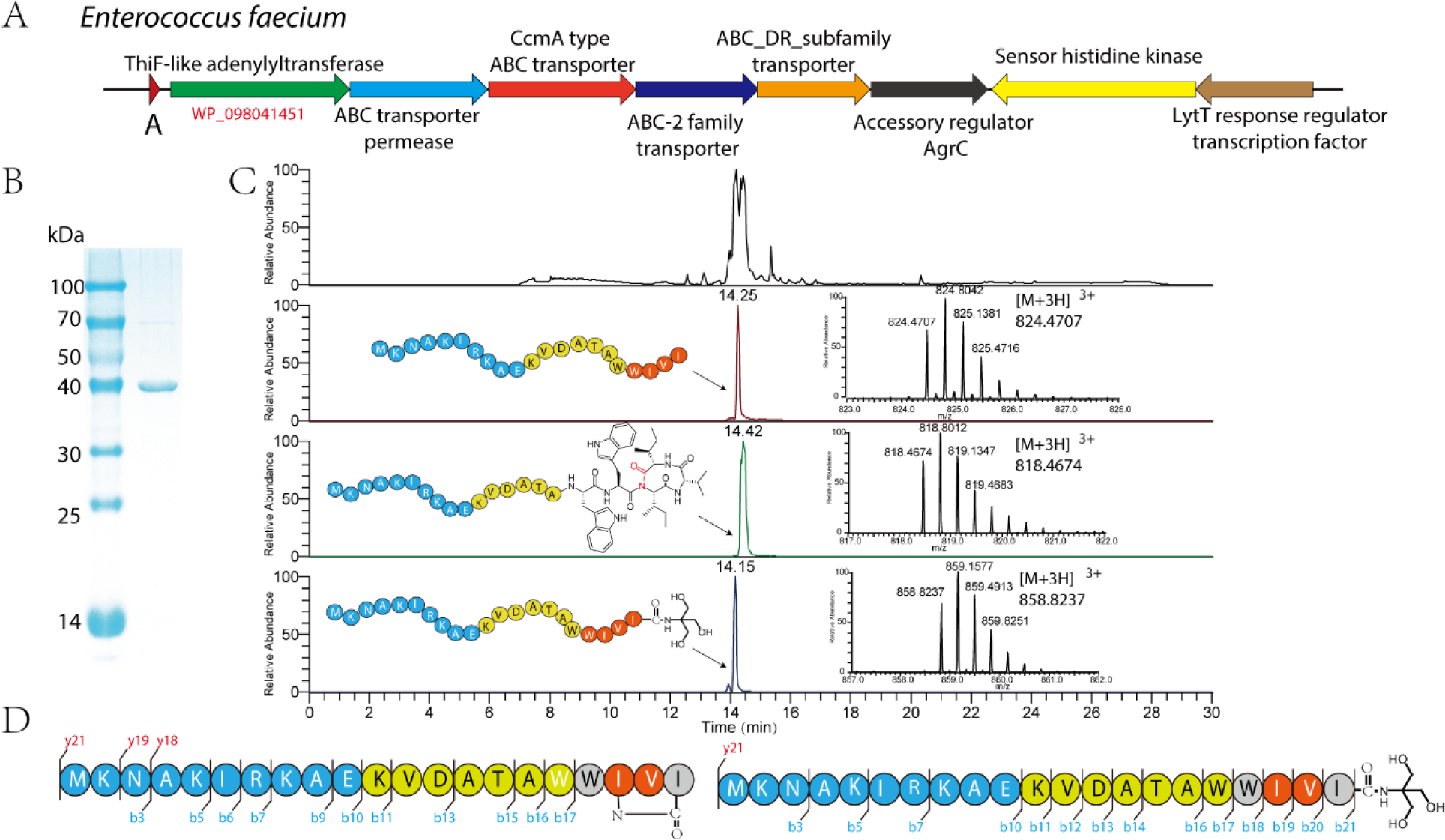
(A) The WWIII cluster identified in *E. faecalis* FDAARGOS_397(WWIVI). (B) SDS-PAGE of the purified EnfB. Lane 1, protein marker; lane 2, the final purified EnfB after size exclusion chromatography. (C) In vitro cyclization and amidation of EnfA precursor peptide by EnfB, as monitored by LC-MS. Extracted ion chromatograms showing the [M+3H]^3+^ ion (m/z = 824.4707) of the precursor peptide, [M+3H]^3+^ ion (m/z = 858.8237) of the amidated peptide and [M+3H]^3+^ ion (m/z = 818.4674) of the cyclized peptide.

**Figure 4.**
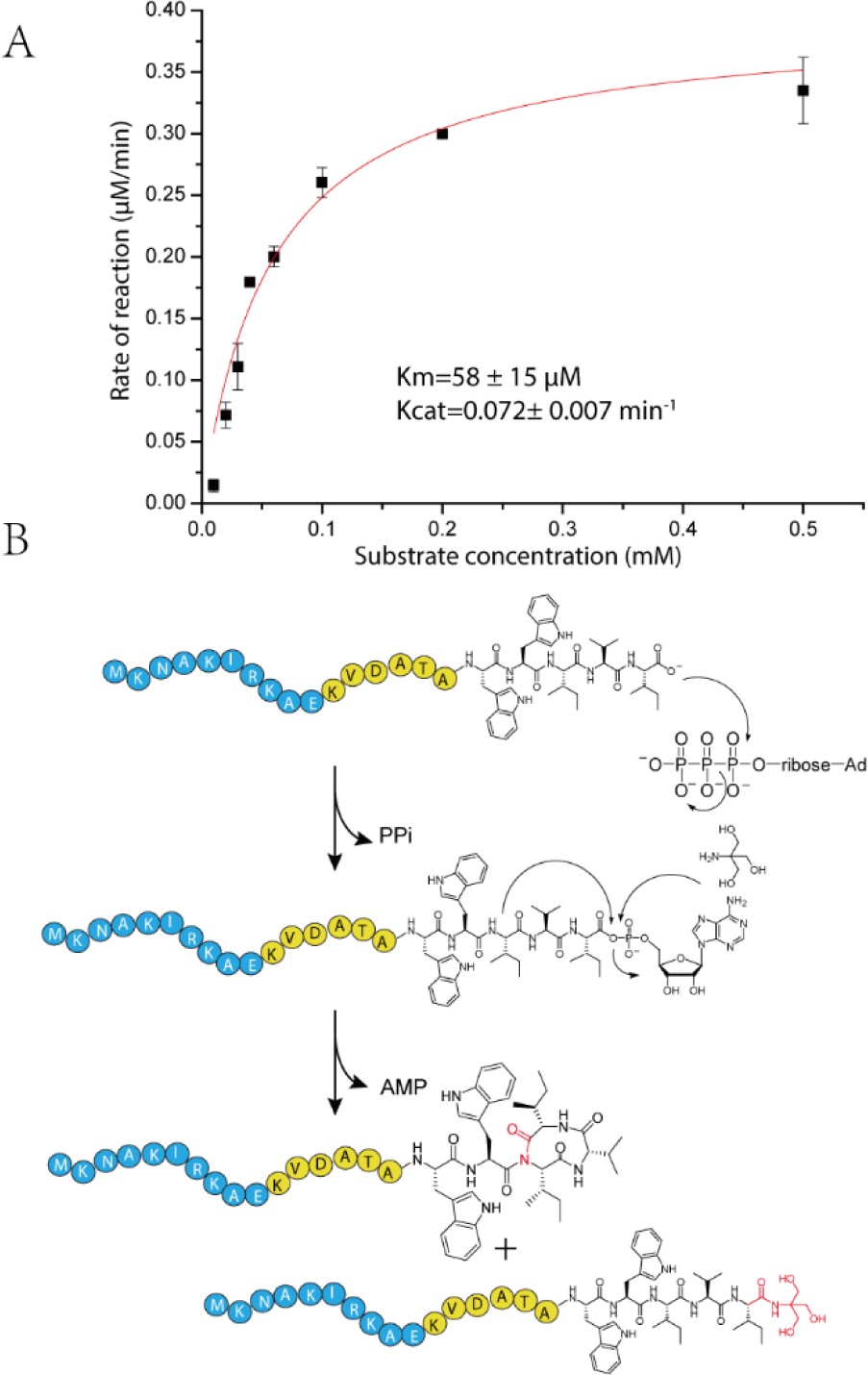
(A) Kinetic characterization of the cyclization catalyzed by EnfB. (B) proposed mechanism of EnfB-catalyzed cyclization and amidation of the C-terminal of a precursor peptide. After activation by ATP, the C-terminal acyl-AMP anhydride of EnfA is either attacked by the nitrogen from the peptide bond at Ile19 to form a macrocyclic imide or by the nitrogen in small organic molecules to amidate the C-terminus.

**Figure 5.**
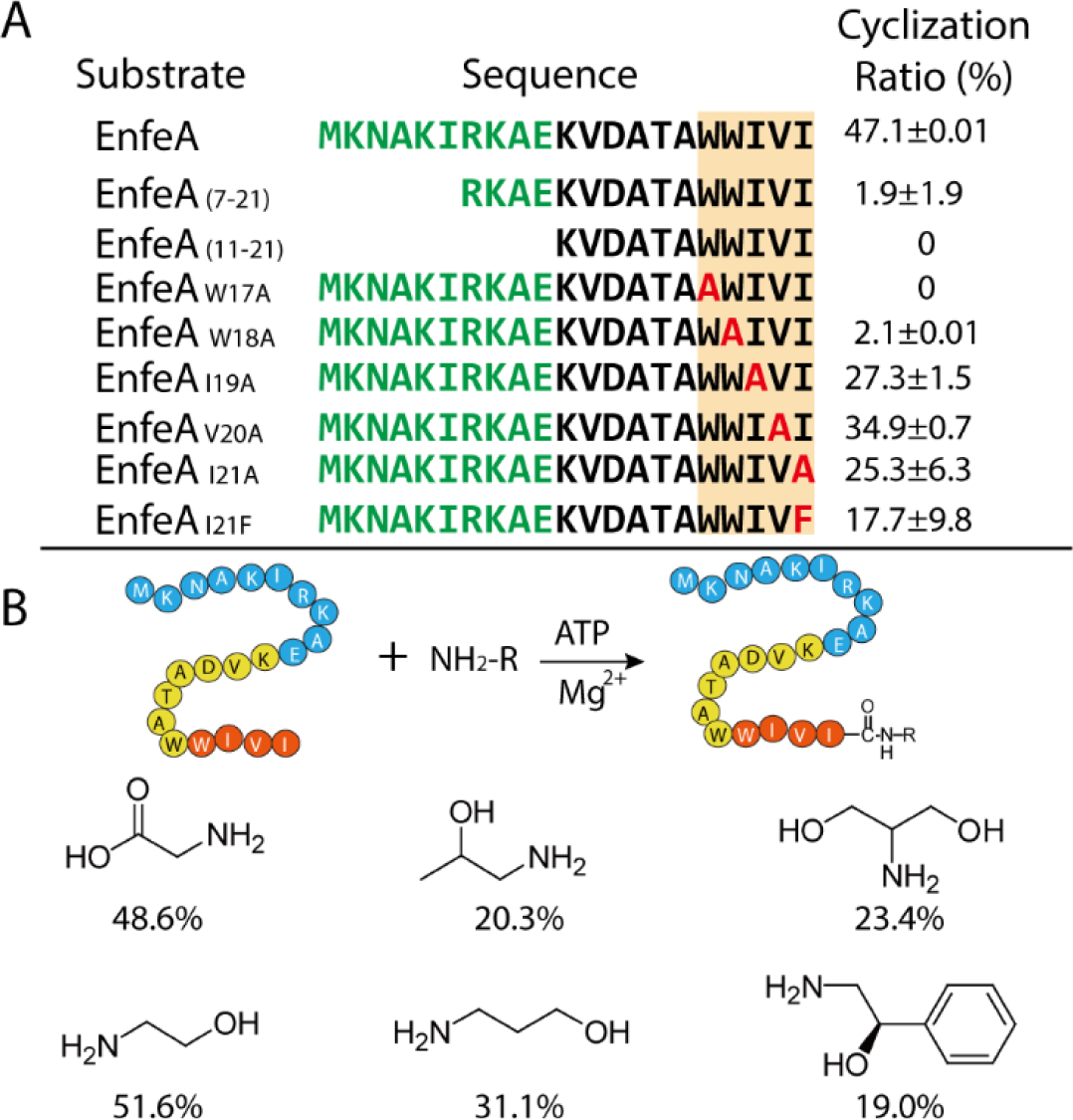
(A) Results of substrate specificity assays with EnfB and EnfA analogues. (B) Results of substrate specificity assays with EnfB and amino compounds. The cyclization ratio or the amidation ratio was shown alongside.

All experiments were conducted in triplicate.

### LC-MS analysis

Mass spectrometric analysis of the *in vitro* studies was conducted by LC-MS. For the structural analysis of the precursor and reaction products, collision-induced dissociation (CID) fragmentation was carried out using online LC-MS. To reveal the cyclization site, a Thermo Scientific Q Exactive Mass Spectrometer connected to a Thermo-Dionex (Ultimate 3000) HPLC system was employed for further MS/MS analysis. Substrates and products were separated on a homemade fused silica capillary column (75 µm ID, 150 mm length; Upchurch Scientific, USA) packed with C-18 resin (300 Å, 5 µm; Varian, USA) using an isocratic gradient of 20% solvent A (0.1% formic acid in water) and 80% solvent B (0.1% formic acid in acetonitrile) at a flow rate of 0.3 mL/min. XCalibur 2.1.2 software was used to control the MS instrument. A single full-scan MS in the orbitrap (100-1800 m/z, 70,000 resolution) followed by 20 data-dependent MS/MS scans at 27% normalized collision energy (higher-energy collisional dissociation, HCD) were performed. Target precursor and enzymatic products were identified and analyzed.

For LC-MS/MS analysis, samples were subjected to a Thermo-Dionex Ultimate 3000 HPLC system, which was directly interfaced with the Thermo Orbitrap Fusion Mass Spectrometer. Peptides were separated on a homemade fused silica capillary column (75 μm ID, 150 mm length; Upchurch Scientific, USA) packed with C18 resin (300 A, 5 μm; Varian, USA) using an isocratic gradient of solvent A (20%, 0.1% formic acid in water) and solvent B (80%, 0.1% formic acid in acetonitrile) at a flow rate of 0.3 μL/min over 120 min. The Orbitrap Fusion mass spectrometer was operated in data-dependent acquisition mode using XCalibur3.0 software, allowing for a single full-scan mass spectrum in the Orbitrap (350-1550 m/z, 120,000 resolution) followed by 3 s data-dependent MS/MS scans in an Ion Routing Multipole at 30% normalized collision energy (HCD).

### Isolation and NMR structure elucidation of the EnfB reaction products

To isolate products of the EnfB reaction, *in vitro* assays were scaled up to 112 mL. A typical scaled-up reaction contained 56 mg EnfA and 5 μM EnfB in HEPES Buffer (50 mM HEPES, 100 mM NaCl, 5 mM ATP, 10 mM MgCl_2_) and was carried out for 3 h at 32 °C with shaking (800 rpm). Reactions were centrifuged and the supernatants were subjected to preparative HPLC (controller, Shimadzu CMB-20A; diode array detector; Shimadzu SPD-20A; pumps, Shimadzu LC-6AD) on a preparative reverse-phase column (SunFire C18 OBD Prep Column, 100Å, 5 µm, 19 mm × 250 mm; Waters) operating at 6 mL/min, with the same gradient as for small-scale reactions. Finally, 5 mg pure product was obtained as white powder and subjected to NMR analysis using DMSO as the solvent.

NMR structural studies on enterofaecin were performed at the Beijing NMR Center and the NMR facility of the National Center for Protein Sciences at Peking University. All spectra were acquired with a 950 MHz Bruker NMR spectrometer. ^1^H−^1^H TOCSY (60 ms mixing time) and ^1^H−^1^H NOESY (150 ms mixing time) experiments were performed. The chemical shift of each proton was assigned based on intra-residue (COSY, TOCSY, HSQC) or inter-residue (NOESY) connectivities.

### Structural modeling of EnfB

To get more structural information about EnfB, AlphaFold Colab (https://colab.research.google.com/github/deepmind/alphafold/blob/main/notebooks/ AlphaFold.ipynb) which is a simplified version of AlphaFold v2.1.0 developed by DeepMind was selected to build the homology modeling structure of EnfB. The AlphaFold modeling studies showed that EnfB has a similar overall structure with the structure of MccB bacterial ancestor of ubiquitin E1 (6OM4) which initiates the biosynthesis of the microcin C7 antibiotic.

## RESULTS AND DISCUSSION

### TLAT-encoding biosynthetic pathways are widespread in lactic acid bacterial genomes

Because RiPP natural products are assembled through a common biosynthetic logic based on conserved features,*^41^* genome mining is an excellent approach to the discovery of previously undescribed compounds. For example, past successes targeting precursor peptides, the radical SAM or YcaO enzyme superfamilies, and even TLATs revealed that myriad “cryptic” RiPPs are yet to be discovered.*^30, 42–44^*

In spite of their fairly small (∼2 Mbp) genomes,*^36^* we hypothesized that of lactic acid bacteria rely extensively on metabolites like bacteriocins to challenge competing organisms. To examine this hypothesis, we focused on TLATs, which, as demonstrated for the RiPPs microcin C and pantocin A, are closely related yet versatile in their biochemical capacities.*^19, 24^* We suspected that mining genomes for TLATs could lead to the discovery of new RiPPs, and thus carried out PSI-BLAST searches against the sequence of MccB from the *E. coli* microcin C system (NCBI code AAY68495.1), with the organism limited to *Enterococcus* (taxid 1350) or *Streptococcus* (taxid 1301).*^37^* Next, 1000 hits from each search were grouped with the EFI-EST web tool and visualized in Cytoscape.*^38, 45, 46^*

In the resulting sequence similarity networks, hits grouped into nine (*Enterococcus*) or eleven (*Streptococcus*) distinct clades (Figs. S1-S2).*^47^* Closer inspection of the TLAT-encoding genes and their genomic surroundings suggested they might have functions in RiPP biosynthetic pathways, as genes directly adjacent code for a putative transporter and a reasonable precursor peptide (Figs. S1-S2). In several cases, a peptidase, transcriptional regulator, and one or more tailoring enzymes (e.g. decarboxylases, methyltransferases, *N*-acetyltransferases, nucleotidyltransferases, PLP-dependent transferases, and radical SAM enzymes) are also encoded. Interestingly, transposons flank a significant number of these putative gene clusters, implying they may have been acquired through horizontal transfer. The genomes of some species, such as *Streptococcus pneumoniae* and *Enterococcus faecalis*, were found to harbor as many as seven potential TLAT-RiPP pathways, supporting our hypothesis that lactic acid bacteria indeed make frequent use of RiPPs type metabolites. Our findings suggest that TLAT-RiPP systems are more complex than previously anticipated and hint at a fertile landscape from which new natural products may be unearthed.

### Diverse TLAT-encoding biosynthetic pathways are found across the bacterial kingdom

We selected six groups of TLATs identified above and investigated their occurrence in all available bacterial genomes beyond those of just lactic acid bacteria. After combining the six groups with TLATs from the microcin C and analogue systems, we constructed a new sequence similarity network (Fig. 2A, Table S1). Notably, TLATs from the microcin C and analogue systems grouped together with no relative from the other six clades. As before, the genomic neighborhood of each TLAT-encoding gene was inspected and annotated with consensus sequences generated by Geneious (Fig. 2B).*^40^* That none of the putative precursor peptides consistently bear a C-terminal Asn residue as in the microcin C system or a central EEN motif as in the pantocin A system suggests they undergo modification disparate from previously reported ones.*^19, 24^*

Second largest among the clades identified (with 108 hits), the WWIII type (named according to the precursor peptide’s C-terminal consensus sequence, WWIVI is this study) is widely distributed among Bacillota (e.g. classes Clostridia and Bacilli, the latter of which includes lactic acid bacteria) in addition to Actinomycetota (Table S1). A diverse set of transporters is encoded by gene clusters from this clade, which is unusual given that natural product biosynthetic pathways typically rely on a single transporter to export a single product; these systems may, therefore, assemble multiple products.*^41, 48^* Another interesting clade closely related to the WWIII type, the NWYFI type, was identified only in lactic acid bacteria (predominantly *Streptococcus* species), with 31 hits. Precursor peptides from both types favor a Trp residue at the preantepenultimate position; however, those from the NWYFI type are nearby double in length. The latter also encode an insulinase-type Zn-dependent peptidase (possibly for leader peptide removal), a transporter, and a transcriptional regulator. The smallest of the identified clades, the VEGR type, is likewise closely related to the WWIII type, though its precursor peptides strictly end in a C-terminal Arg residue. In the biosynthesis of microcin C, the side chain of the C-terminal Asn residue is crucial for N-P bond formation;*^24^* the conserved Arg residue in the VEGR type may be similarly important for post-translational modification, though this hypothesis awaits experimental validation. Eight of the nine hits from this clade come from Bacillota (class Bacilli), with the last one from Fusobacteria.

Two relatively small clades (with 23 and 24 hits, respectively), the NICxxxK and CxxxC types, were found almost exclusively in *Streptococcus* genomes. A TLAT-peptidase fusion is encoded by gene clusters of the latter type. In both types, regulators are encoded, and the strictly conserved C-terminal residue may participate in post-translational modification, as with the VEGR type. Finally, the largest clade (with 131 hits), the GxxQY type, is the only to code for two distinct TLATs. Aside from Bacillota, hits from the GxxQY type appear extensively in Proteobacteria (including human pathogens like *Salmonella enterica*, *Acinetobacter baumannii*, and *Klebsiella pneumoniae*) with additional members from diverse phyla including Aquificota, Bacteriodota, Campylobacterota, Fusobacteria, and Spirochaetota. Based on the organizations, compositions, and precursors of the abovementioned gene clusters, our bioinformatics analyses suggest nature employs TLATs extensively to supplement the functionality of RiPPs.

### Characterization of a WWIII-type gene cluster

To verify that TLATs in the identified RiPP systems can catalyze new biotransformations, we chose a WWIII-type gene cluster for further investigation. This type is dominant in *Enterococcus* and *Streptococcus* species – notably, *E. faecalis* and *E. faecium*, which are commensals and opportunistic pathogens in the gastrointestinal tracts of humans,*^34, 49, 50^* and *S. thermophilus*, which is exploited in the production of yogurt.*^51^* Here, we focused on gene clusters from *E. faecalis* FDAARGOS_397, a species notorious for its multidrug resistance.*^50, 52^* The WWIII-type gene cluster encodes a 21-residue precursor peptide (EnfA, which has a WWIVI motif in this study), a TLAT (EnfB), four transporters (EnfC-F), and three enzymes involved in regulation (EnfG-I) (Table S3, Fig. 3A). Due to the lack of a leader peptidase in the gene cluster, we assumed the final product requires its leader sequence for activity and/or cellular uptake, similar to microcin C.*^27, 53^* It is also possible that a potential peptidase located in another region of the genome could be responsible for the removal of the leader peptide. However, this hypothesis needs to be confirmed in future research.

To assess the TLAT reaction, we purchased the precursor peptide EnfA prepared by solid-phase peptide synthesis; the TLAT EnfB was heterologously expressed in *E. coli* with a C-terminal His_6_ tag and purified via Ni-NTA and size-exclusion chromatography (Fig. S3, Fig. 3B). EnfA and EnfB were then incubated together in the presence of an ATP, CTP, UTP, or GTP cosubstrate in Tris buffer and monitored by HPLC. While no change was observed in reactions containing CTP or GTP, two new peptide species appeared when ATP or UTP was included (although efficiency was higher for ATP) (Fig. S4). This observation resembles that made with MccB from *E. coli* and *B. amyloliquefaciens*, which favor ATP and CTP, respectively, but are able to modify their precursor peptide in the presence of any of the four NTPs.*^28^* In the biosynthesis of microcin C, a peptidyl-succinimide intermediate was observed.*^24^* We also anticipated formation of an intermediate in our reaction, which would explain the presence of two new species.

### MS and NMR analyses reveal a novel cyclization-type post-translational modification

To determine the identity of the peptide products observed in our EnfB activity assays, we carried out high-resolution MS/MS analysis. Relative to native EnfA ([M+3H]^3+^ calc: 824.4701, [M+3H]^3+^ obs: 824.4707), one product showed loss of water ([M+3H]^3+^ obs: 818.4674, [M+3H]^3+^ calc: 818.4666), suggesting EnfB promotes intramolecular cyclization of the peptide (Fig. 3C, Fig. S5).*^24^* Although we could observe most of the b-ion fragments for the substrate, fragmentation was not detected within the final four residues of the product, pointing to one of those residues and the terminal carboxyl group as the likely sites of cyclization (Fig. 3D, Fig. S6). The second product was differently modified – not with an adenylyl group but a Tris group ([M+3H]^3+^ obs: 858.8237, [M+3H]^3+^ calc: 858.8245). Indeed, when Tris was omitted from the assays, the modification was absent from EnfA (Fig. S7). Observation of the b21 fragment pointed to the terminal carboxyl group as the likely site of modification (Fig. S6).

To confirm our hypothesis, further LC-MS/MS analysis was performed and a similar result was obtained. The observation of the b18 ions in both native EnfA and Tris-modified EnfA and the absence of b18 ions in the cyclic product are consistent with the previous observation, which provided a fragmentation pattern indicative of cross-link formation at the C-terminal WIVI motif.

To further pinpoint the position of EnfB-catalyzed cyclization, structure elucidation was conducted via NMR spectroscopy. Scaling up of *in vitro* assays and purification by preparative HPLC resulted in 2 mg cyclized peptides (Fig. S8), which were subjected to extensive NMR analysis (Fig. S9-Fig. S14). We were able to completely assign resonances to the linear EnfA substrate (Table S4); in contrast, assignment of the product resonances was more challenging due to spontaneous hydrolysis. Nevertheless, compared to linear EnfA, several pieces of evidence suggest formation of a macrocyclic imide at the C-terminal motif. In the substrate spectra, two typical indole signals (10.24 and 10.04 ppm) were apparent for Trp17 and Trp18. In the product spectra, these signals were likewise present, which ruled out the possibility of an isopeptide bond involving the indole amine of either Trp side chain and instead suggested an imide bond between the Ile19 backbone N atom and the C-terminus. These results, combined with our MS analysis, suggest EnfB installs macrocyclic imides in peptides but also activates their C-termini for modification. However, since the NMR analysis of the cyclization products has not been fully resolved, further research is necessary to confirm their detailed structural information.

### Biochemical characterization of EnfB

EnfB-catalyzed conversion of EnfA was subsequently characterized *in vitro* through time course analysis in phosphate buffer by HPLC After 6 h, ∼50% of the precursor (200 μM) was transformed into a macrolactam by 1.75 μM EnfB (Fig. S15). Increasing the reaction duration failed to improve conversion significantly, likely due to enzyme instability (a white precipitate became visible during the assays). Therefore, incubation for 3 h was selected for further optimization. Interestingly, we observed a steady increase in EnfB activity with temperature from 10 to 50 °C, with a decline to nil by 60 °C. EnfB thus exhibits modest thermostability and processes its substrate most efficiently at a temperature significantly higher than that for optimal growth of *E. faecalis* (Fig. S16A). We also tested the effect of pH on EnfB activity over a range of 4.0 to 10.0 (at 50 °C), resulting in a maximum at pH 7.0 (Fig. S16B). Finally, the kinetics with respect to EnfA concentration were evaluated at the optimal pH and temperature, resulting in apparent *K*_m_ = 58 ± 15 μM, *k*_cat_ = 0.072± 0.007 min^-1^, and *k*_cat_/*K*_m_ = 0.001 min^-1^ μM^-1^ (Fig. 4A).

### Proposed catalytic mechanism of EnfB

Based on the consumption of ATP and formation of two products in our assays with EnfB, the reaction likely resembles that of MccB, which catalyzes C-terminal adenylation of its precursor peptide followed by intramolecular cyclization and subsequent ring opening.*^24^* To compare the structures of these two enzymes, (EnfB 22% identical in sequence to MccB), we predicted its structure with the neural network AlphaFold (Fig. S17).*^54, 55^*

As observed for MccB and PaaA, the EnfB model consists of two subdomains: an N-terminal RIPP recognition element (RRE) domain and a C-terminal adenylation domain that bind and modify the precursor peptide leader and core sequences, respectively.*^11, 19^* Superposition of the EnfB model with the structure of MccB bound to ATP and Mg^2+^ (PDB ID: 6OM4) allowed the roles of key residues to be predicted. Arg167 and Lys180 are highly conserved among reported TLATs due to their essential role in ATP binding.*^11, 19, 56^* When we substituted these residues with Ala, EnfB was found to be insoluble. Asp225, another conserved residue, is expected to coordinate the Mg^2+^ ion that stabilizes the negative charge of ATP. Ala mutagenesis likewise showed this residue is essential, as no conversion of EnfA was detected. Moreover, an ordered ‘lid’ motif, not modeled in the MccB structure, was observed above the putative substrate binding pocket.

On the basis of the structural information, we proposed EnfB employed a similar catalytic mechanism for C-terminal carboxyl activation of EnfA but a different process for further post-translational modification. While MccB requires two moles of ATP for N-P bond formation, EnfB only consumes one mole of ATP for cyclization and amidation.*^24^* The first stage is similar, whereas the C terminus carboxylate attacks P_α_ of the ATP, yielding a hydrolytically labile precursor-CO-AMP. In the modification of MccA, the carboxamido nitrogen of Asn7 first attacks the C-terminal acyl-AMP anhydride to form a succinimide intermediate, then a second ATP is consumed to form the unusual N-P bond to synthesize the unprocessed microcin C. On the other hand, there is a different scenario for enterofaecin biosynthesis. The C-terminal acyl-AMP anhydride of EnfA was either attacked by the nitrogen from peptide bond at Ile19 to form a macrocyclic imide or by the nitrogen containing small organic molecules to amidate the C terminus (Fig. 4B).

### Substrate specificity of EnfB

We next investigated the substrate specificity of EnfB. Given that RiPP biosynthesis is guided by the leader sequence, we wondered whether the enterofaecin system would comply with the same rules.*^48^* Two N-terminal truncations, EnfA_(11-21)_ and EnfA_(7-21)_, which lack a conserved (V/I)RKA motif in the leader sequence and all residues N-terminal to that motif, respectively, were prepared (Figs. 2A and 5A). Following incubation with EnfB for 3 h, no or minor conversion was observed by LC-MS analysis, in agreement with the role of the putative N-terminal RRE domain of EnfB in binding the leader sequence.

We next probed the roles of Trp17 and Trp18 in the core sequence, which are conserved in many of the identified gene clusters of the WWIII type. Substitution of Trp17 or Trp18 with Ala completely abolished conversion or decreased it to 2.1%, respectively, suggesting these residues are likely involved in substrate recognition by the adenylation domain of EnfB. We considered whether substitution of residues in the ring structure would disrupt enzymatic processing. The variants EnfA_I19A_, EnfA_V20A_, EnfA_I21A_, and EnfA_I21F_ were generated and individually incubated with EnfB as before. As yields ranged from 17.7−34.9% for these variants, EnfB is apparently relaxed in specificity towards the three C-terminal positions of EnfA (Fig. 5A; Figs. S18-S21).

As noted above, we observed C-terminal amidation of EnfA with Tris when assays with EnfB were carried out in Tris buffer. To further probe the substrate scope of the enzyme, we assayed a small set of primary amines: glycine (**1**), isopropanolamine (**2**), serinol (**3**), ethanolamine (**4**), propanolamine (**5**), and (*R*)-phenylethanolamine (**6**) (Fig. 5B). Interestingly, EnfB was able to catalyze the amidation of EnfA with all six of these compounds (Fig. 5B, Fig. S22-S27). Coupled with the presence of a multitude of primary amines in the cell, this activity may explain why multiple transporters are encoded in the enterofaecin gene cluster.

### Evolution of isopeptide macrocyclase

Our genome mining efforts revealed that TLAT-RiPP pathways are widely present in nature. For example, the WWIII type is utilized by a range of Bacillota (including lactic acid bacteria) and Actinomycetota. To study the evolutionary patterns of this type of system, we prepared a phylogenetic distribution of precursor peptides through the neighbor-joining method.*^57^* As demonstrated in Fig. 6A, precursor peptides from different phyla group into several subclades. Closer examination of the encoding gene clusters suggests different architectures for different precursor peptides (Fig. 6B). We generally classify these systems into 4 classes based on their biosynthetic pathways. Type I, as unveiled in *E. faecium*, features up to 4 different transporters and is highly conserved as WWxxx, though several RWxxx cases were observed. Type II consists of precursors highly conserved as SWxxx, which are mainly synthesized from gene clusters featuring only two transporters. Both systems are closely related to a LytT response regulator transcription factor and a sensor histidine kinase, suggesting they perform a similar function. On the other hand, type III is coevolved with an MFS transporter, a CitB response regulator transcription factor and a 2CS histidine protein kinase, suggesting a completely different function. The core sequence of the precursor diversified during evolution; however, relatively small hydrophobic amino acids such as Ile, Leu, Val and Gly are highly preferred. In several small groups, Phe is also preferred indicating the broad substrate specificity of the ThiF enzymes and the structural diversity of the natural products. These bioinformatic results also are consistent with the in vitro studies on the substrate specificity of EnfB. Notably, type IV system from *Nonomuraea* sp. PA05 has evolved independently with a NWVFF core, supporting the idea that substantial diversity of macrolactam has been developed in nature. More rigorous bioinformatics analyses in future studies might uncover types with distinct sequence and properties.

**Figure 6.**
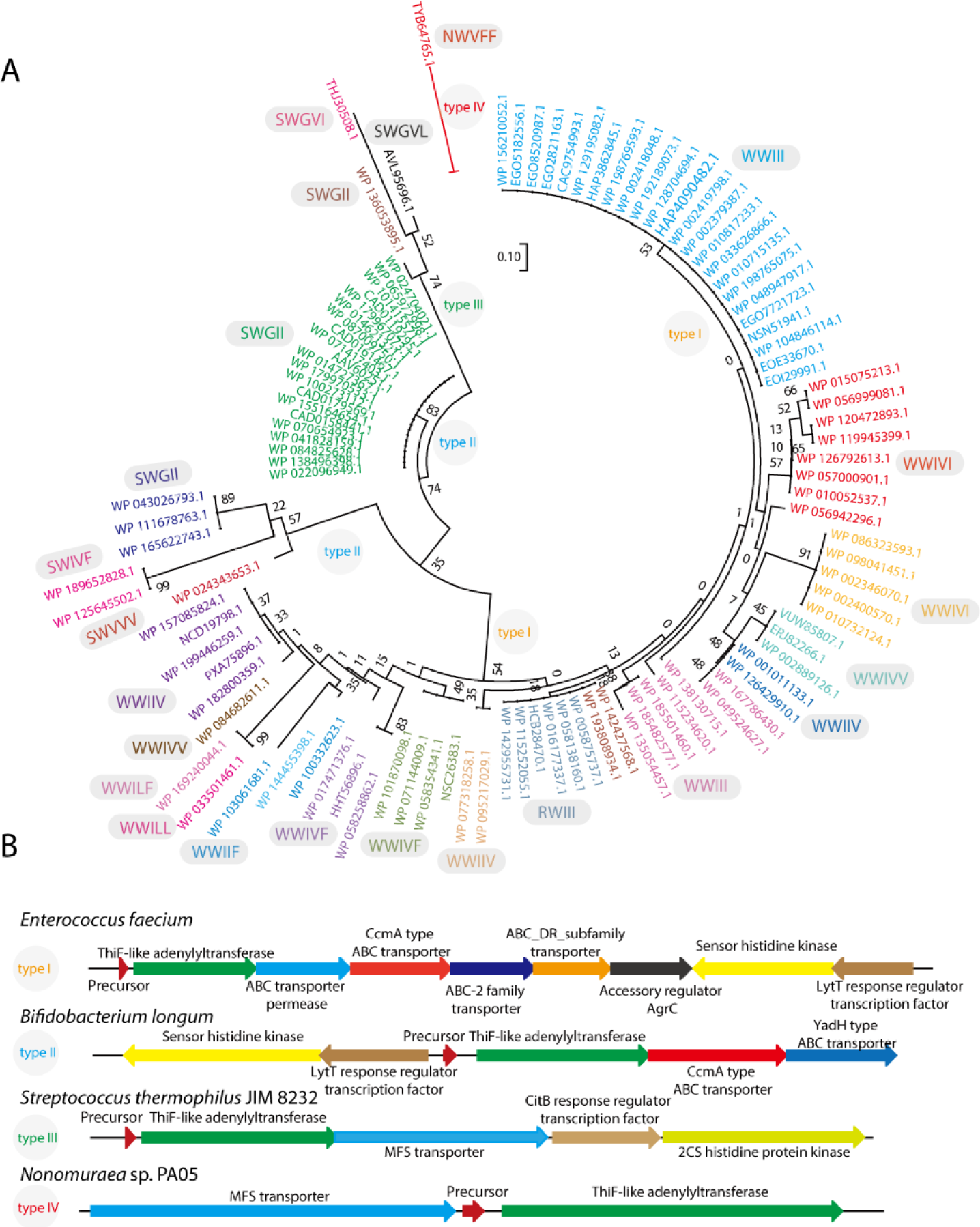
(A) Evolution of the precursor peptide for these diverse macrocyclase. The substrates could be classified into 4 clades based on the organization of their biosynthetic gene clusters. (B) Representative biosynthetic gene cluster for each of the sub - families in panel A.

Taken together, these results indicate that these systems from different phyla have evolved independently with potential different functions. However, the XWXXX motif is strictly conserved, suggesting a similar macrocyclic imide has been developed, though further experimental assessment is need. The evolution of these macrocyclases also greatly enrich our toolkits for construction of new cyclic framework.

### Bioactivity of enterofaecin

Also unknown is the function of these macrocyclic imides. In history, lactam and macrolactam compounds have been the mainstays of antibiotic therapy. Therefore, investigation of the biological role of these macrolactam peptides is of great interest.*^58^* Notably, downstream of these type gene cluster are three genes including an accessory regulator AgrC domain, a sensor histidine kinase, and a LytT response regulator transcription factor, which is easily reminiscent of the QS accessory gene regulator (Agr) locus system in *Staphylococcus aureus*.*^59^* QS are key intercellular signaling process that are essential for many bacteria to control their population densities and virulence induction.*^60, 61^* Many pathogenic bacteria utilize QS to launch synchronized attacks on their hosts only after they have achieved a high cell density, thereby overwhelming the host’s defense mechanisms.*^61^* In *S. aureus*, the QS-Agr system encodes four genes, which produce a secreted autoinducer peptide (AIP) and receptor histidine kinase AgrC (Fig. S28).*^62–64^* The membrane protease AgrB is required for processing of AgrD into AIP, a macrocyclic peptide (7–9 residues) containing an N-terminal exocyclic tail and a thiolactone bridge between an internal Cys side chain and the C-terminus.*^65^* When a threshold extracellular AIP concentration is reached, cognate AIP-AgrC interactions then act to phosphorylate the LytT transcription factor. The phosphate LytT then binds to several promoters to autoinduce the agr system and upregulate RNAIII transcription, by which controlling over 100 virulence factors (Fig. S28).*^63, 64^* Compared with different types of AIP and enterofaecin, the five residue macrocyclic thiolactone core in AIP is similar with the macrolactam in enterofaecin regarding primary sequences which mainly consist of several small hydrophobic amino acids (Fig. S28). To study the function of this gene cluster, a structural model of the sensor histidine kinase was generated (Fig. S29).

As predicted, the sensor histidine kinase could be divided into a transmembrane domain and a C terminal kinase domain indicating it is indeed a QS system. Notably, while these gene clusters are widely present in different *E. faecium* strains, several strains only have a truncated version of the EnfB gene (Fig. S30). We bought both the *E. faecium* strain ATCC 51299 (full length EnfB homology) and *E. faecium* strain 15244 (truncated EnfB homology). Fermentation of each strain resulted in a different growth curve. As shown in Fig. S31, *E. faecium* strain ATCC 51299 has an obvious QS system that regulates cell density after 20 hours, with a maximal OD of about 1.5. In contrast, the OD of *E. faecium* strain 15244 can reach 4.0 after 30 hours. This suggests that these two strains have different QS systems, which may be due to the inactive EnfB in *E. faecium* strain 15244. It is also possible that another QS system may be responsible for the different growth curves. However, the association of an obvious QS system with enterofaecin gives us a strong hint that might be a novel QS signal. Further studies are necessary to demonstrate our hypothesis.

## CONCLUSION

Microbial natural products that have progressed to clinical utility in humans are largely of polyketide or peptide origin initially synthesized as a linear precursor or acyclic nascent chains tethered at the termini of their constituent NRPS and PKS assembly lines, which need undergo further enzymatic morphing to generate bioactive framework complexity.*^58, 66^* A majority of the mature active products frequently have developed constrained scaffolds via macrocyclizations or heterocyclizations that endow them with notable chemical stability and conformation for targets.*^4, 58, 67^* Therefore, development of novel chemo-enzymatic route to macrocycle formation in natural product biosynthesis is of great interest. RiPPs are a rapidly growing family of natural products with mechanistically distinct logic and enzymatic machinery for cyclization. The subcategories of RiPPs have expanded greatly due to the explosion of genomic data and the development of novel bioinformatic approaches. While past researches have intensively focus on talent genus for natural products discovery, bacterial genera with small genomes such as lactobacillales have been frequently overlooked.*^68^* We herein significantly expand the ThiF-RiPPs family of natural products by genome mining in lactobacillales and systematically identified diverse ThiF-RiPPs groups based on the type of biosynthetic pathways and the conserved precursor sequences. This bioinformatics survey based on sequence similarity network limited one genus proved phylogenetically distinct but unattended subgroup of biosynthetic machineries might be rich sources for novel natural products with completely new scaffolds. Focusing one system from *E. faecalis* FDAARGOS_397 identified in the network, we demonstrated a distinct logic to form constraining macrocyclic imides in diverse peptides. This discovery provides a remarkably efficient strategy to assembly peptide into highly compact macrocyclic framework. Previously, peptide macrolactam framework are frequently synthesized via giant NRPs machinery which use the thioesterase domains to mediate macrocyclic formation.*^66^* Macrolactam construction mediated by ATP are found in only a few RiPP families, such as lasso peptides and ω-ester-containing peptides.*^69, 70^* However, the reaction of EnfB here described is the first imide macrocyclase that specifically catalyzes the cyclization of its precursor peptide by connecting a conserved WI(/V)V(/I)I(/V) motif in the C terminus by macrocyclic imide. Considering the fact that lactams type natural products are still the mainstays of antibiotic therapy, this distinct biochemical logic and post-translational enzymatic machinery is of great use to create diverse arrays of macrocyclic imide structures.

In conclusion, this paper lays the foundation for further study of novel ThiF-RiPPs type natural product biosynthesis, in which many intriguing questions remain. The WWIII system described here produces both cyclic and C-terminus amidated products that likely require different types of transporters for export. Thus far, the function of these peptide molecules is still unknown. The association of accessory regulator AgrC, the sensor histidine kinase, and the LytT response regulator transcription factor genes suggest they might be involved in quorum sensing (Fig. 7). *^60^* Since several prevalent human pathogens (e.g., *S. aureus*) use QS to control virulence, this potential QS system is of great interest, given that diverse *Streptococcus* and *Enterococcus* species are also pathogens. *^60,71^* The QS system might be developed as a novel anti-infective target. The investigation here into the biosynthesis of novel macrolactam analogues thus provides a new opportunity in the development of non-native ligands capable of interacting with such QS pathways.

**Figure 7.**
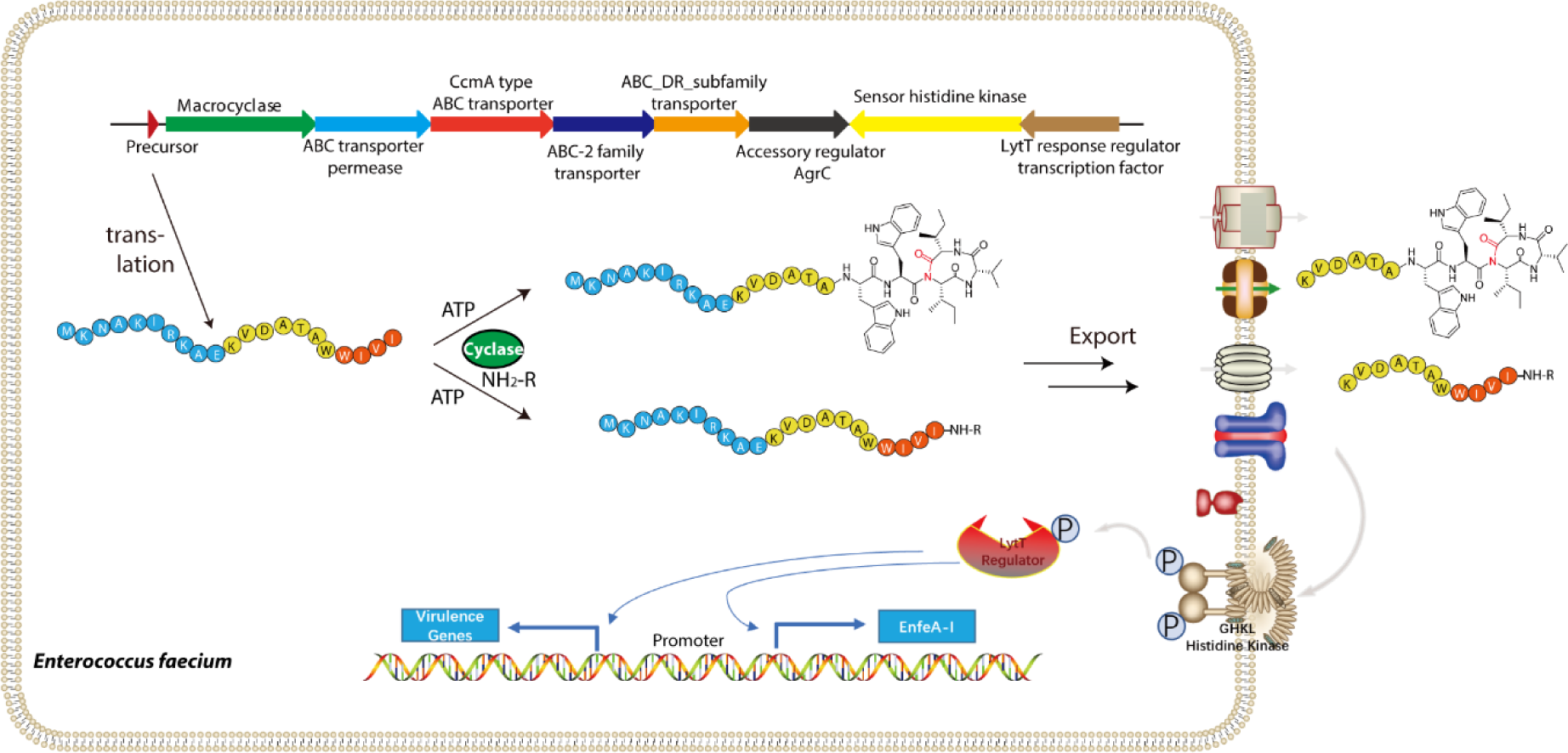
(A) Proposed schematic of enterofaecin biosynthesis. Precursor peptide EnfA is first either cyclized or amidated. A portion of these peptides are then exported by diverse transporter, while triggered the potential QS system.

In addition to the system characterized here, the discovery of diverse ThiF-RiPPs gene clusters in Lactobacillales also requires further exploration to identify the mature pathway end products, which warrant additional investigation regarding their chemical structures and biological functions. We believe that more rigorous bioinformatics analyses will uncover additional novel RiPPs groups with distinct structures and properties, thereby providing insights into biosynthetic mechanisms and other tailoring modifications. Focusing solely on ThiF-RiPPs type natural products in *Streptococcus* and *Enterococcus* has been fruitful, providing novel systems that might only represent the tip of the iceberg. Future investigations into other genera targeting ThiF and other biosynthetic enzymes, such as the YcaO and RaS families, will offer great opportunities for the discovery of novel RiPPs, as well as for understanding the underlying biology, chemistry, and enzymology in the coming years.

## ACKNOWLEDGEMENT

Financial support from Natural Science Foundation of Beijing Municipality (No. 7202107), the National Natural Science Foundation of China (NSFC; Grant No. 21706005) and National Key R&D Program of China (2022YFC2804900) are gratefully acknowledged. All NMR experiments were carried out at the Beijing NMR Center and the NMR facility of National Center for Protein Sciences at Peking University.

## ASSOCIATED CONTENT

Supporting information available: This material is available free of charge via the internet.

### Accession Codes

The *EnfB* gene from *E. faecalis* FDAARGOS_397 strain (Accession number: WP_098041451.1)

## AUTHOR INFORMATION

### Corresponding Authors

*(S.Z.) Tel.: +86-10-64431557.

*(A.F.) Tel.: +86-10-64431557.

*(Y.T.) Tel: +86-10-64431557.

### Author Contributions

M. Wang., and M. Wu contributed equally. Conceptualisation, supervision and writing – S. Z., A. F and Y. T.; investigation and methodology –M. W., M. W., M. H., X. N.; all authors have given approval to the final version of the manuscript.

### Notes

The authors declare no competing financial interest.

## Notes

### Competing Interest Statement

The authors have declared no competing interest.

