## Supplementary data for "Mining the biosynthetic landscape of lactic acid bacteria unearths a new family of RiPPs assembled by a novel type of ThiF-like adenylyltransferases"

**Supporting information**

### Supplementary Figures and Tables

**Supplementary Table S2**. Primers used in this study to prepare variants of EnfB.

| Construct | Primer name | Primer sequence |
|---|---|---|
| R167A | R167A-P1 | TTA TCG CGA GAA TGA TAT TGG AAA AAA GAA AAC |
| | R167A-P3 | ATCTCGGTGCACAA TTT CTT TAT CGC GAG AAT GAT ATT GG |
| | R167A-P2 | TTCCAAGTTCGACAATGTCATAATCGACTA |
| | R167A-P4 | AGAAATTGTGCACC GAG ATT TCC AAG TTC GAC AAT GTC |
| K180A | K180A-P1 | TGC TAA GAG TGC AGT AGA AAA AAT GAA TTC |
| | K180A-P3 | GAAAAAAGGCA ACA GAA ATT GCT AAG AGT GCA GTA GAA AAA AT |
| | K180A-P2 | CAA TAT CAT TCT CGC GAT AAA GAA ATT GTCT |
| | K180A-P4 | ATTTCTGTTGCCTT TTT TCC AAT ATC ATT CTC GCG ATA AAG |
| D225A | D225A-P1 | ATT TAT TAT TCA TCG AAT AGT GAA TGA AGC GAT T |
| | D225A-P3 | GTGCAATTGCAGAG CCT CCA TTT ATT ATT CAT CGA ATA GTG AAT |
| | D225A-P2 | AGA CGA CTA TGT CAA CAT CAT TTA TTA GTT CTA AT |
| | D225A-P4 | GGAGGCTCTGCAAT TGC ACA GAC GAC TAT GTC AAC ATC ATT |

**Supplementary Table S3.** Proposed functions of ORFs in the RiPP BGC from Enterococcus faecium containing a TLAT.

| **Gene product** | **Length (aa)** | **NCBI code** | **Proposed function** | **Conserved domain analysis** |
|---|---|---|---|---|
| EnfA | 21 | - | precursor peptide | - |
| EnfB | 375 | WP_098041451.1 | processing enzyme | RIPP recognition element (RRE) domain, ThiF family, thiazole biosynthesis adenylyltransferase |
| EnfC | 288 | WP_227644887.1 | export, immunity | ABC-type transporter permease |
| EnfD | 308 | WP_098041450.1 | export, immunity | ABC-type transporter, ATPase component (COG1131: CcmA) |
| EnfE | 257 | WP_086310538.1 | export, immunity | ABC-2-type transporter, transmembrane component (pfam12698: ABC2_membrane_3) |
| EnfF | 258 | WP_086323637.1 | export, | ABC_DR_subfamily transporter, ATP binding |

| | | | | |
|---|---|---|---|---|
| | | | immunity | component (cd03230: ABC_DR_subfamily_A) |
| EnfG | 245 | WP_227644886.1 | regulation | Accessory regulator AgrC, AgrC domain-containing protein |
| EnfH | 428 | WP_086323594.1 | regulation | Sensor histidine kinase, histidine kinase-like ATPase domain of two-component sensor histidine kinases, similar to *Staphylococcus aureus* AgrC and *Streptococcus pneumoniae* ComD, which are involved in quorum sensing |
| EnfI | 236 | WP_029682523.1 | regulation | LytT response regulator transcription factor, DNA-binding response regulator, LytR/AlgR family |

**Supplementary Table S4.** Assignment of $^1$H signals (ppm) from linear EnfA (950 MHz, DMSO-$d_6$).

| Residue | NH | Hα | Hβ | Others |
|---|---|---|---|---|
| Met1 | 8.17, d, 7.9 | 4.23, m | 1.94, m | 2.30, m:Hγa 1.80, m:Hε* |
| Lys2 | 8.22, d, 5.2 | 4.18, m | 1.59, m; 1.66, m | 1.53, m:Hγa 1.27, m:Hδa 2.83, m:Hεa |
| Asn3 | 8.46, d, 6.9 | 4.52, m | 2.62, m; 2.70, m | |
| Ala4 | 8.20, d, 6.0 | 4.16, m | 1.26, d, 7.4 | |
| Lys5 | 8.64, d, 6.6 | 4.22, m | 1.63, m; 1.67, m | 1.29, m:Hγa 1.56, m:Hδa 2.86, m:Hεa |
| Ile6 | 7.93, d, 8.4 | 4.02, m | 1.70, m | 1.33,m :Hγ1a 1.05, m:Hγ2* 0.75, m:Hδ1* |
| Arg7 | 8.29, d, 7.2 | 4.22, m | 1.59, m; 1.68, m | 1.44, m:Hγa 1.49, m:Hγb 3.03, m:Hδa 7.04, m:Hη1a |
| Ala9 | 8.23, d, 7.5 | 4.15, m | 1.23, d, 7.5 | |
| Lys11 | 8.19, d, 7.7 | 4.16, m | 1.60, m; 1.67, m | 1.54, m:Hγa 1.32, m:Hδa 2.85, m:Hεa |
| Val12 | 8.06, d, 7.8 | 3.95, m | 1.91, m | 0.78, m:Hγa* |
| Asp13 | 8.38, d, 7.1 | 4.54, m | 2.66, dd, 17.4, 7.3; 2.78, dd, 17.4, 6.2 | |
| Ala14 | 8.15, d, 5.8 | 4.18, m | 1.27, d, 7.4 | |
| Thr15 | 7.85, d, 7.8 | 4.05, m | 3.97, m | 0.98, d, 6.7:Hγ1 |
| Ala16 | 7.82, d, 6.4 | 3.75, m | | |
| Trp17 | 7.45, brd, 5.7 | 4.34, m | 3.07, dd, 15.7, 5.8; 3.19, dd, 15.7, 5.6 | |
| Trp18 | 6.92, brd, 7.3 | 4.37, m | 2.43, m; 2.93, m | |
| Ile19 | 7.28, brd, 7.7 | 3.92, m | 1.54, m | 1.00, m:Hγ1a 0.72, m:Hγ2* 0.63, m:Hδ1* |
| Val20 | 7.93, d, 8.4 | 3.99, m | 1.91, m | |
| Ile21 | 8.07, d, 8.1 | 4.11, m | 1.77, m | 1.31, :Hγ1a 1.08, m:Hγ2* 0.73, m:Hδ1* |

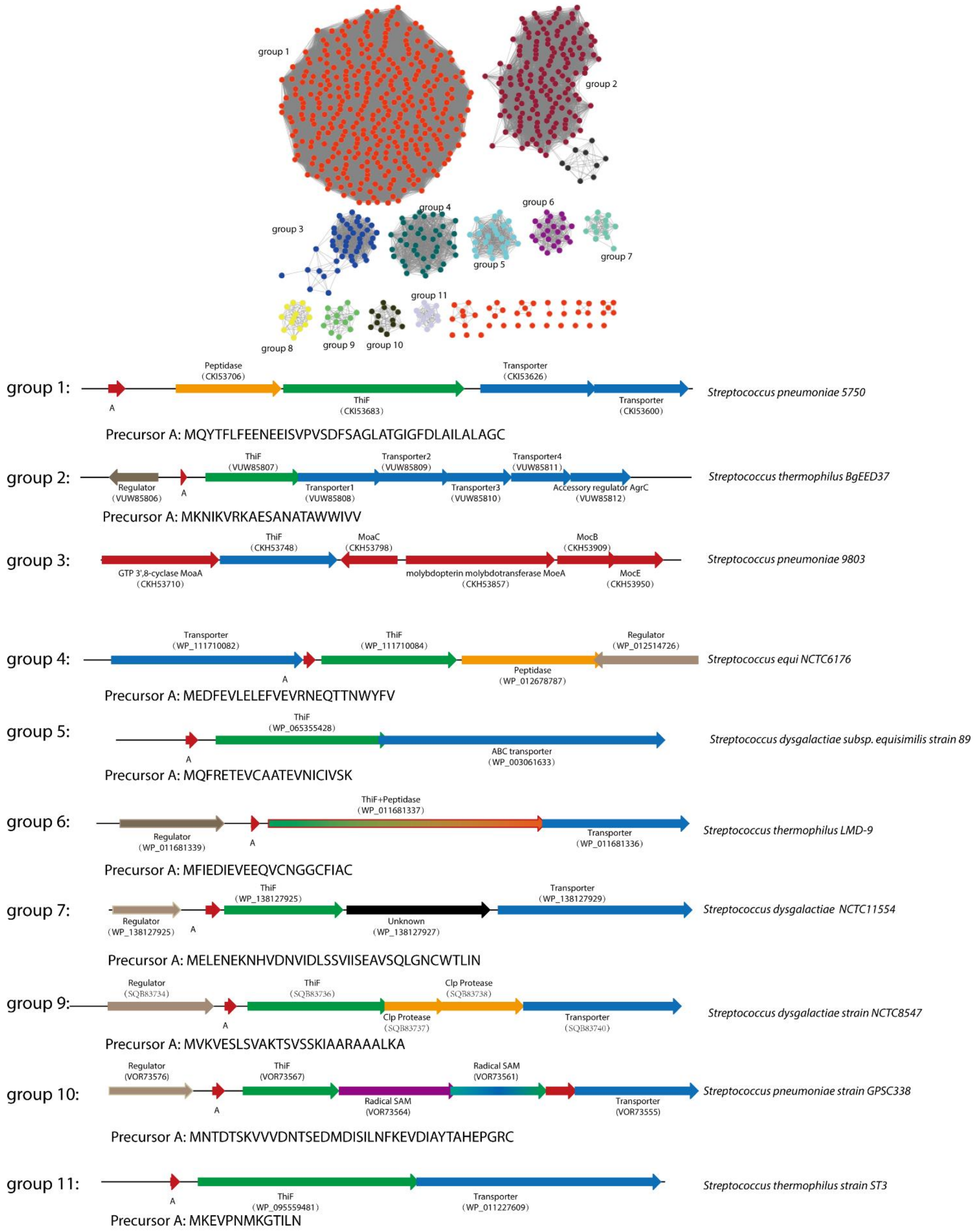


**Supplementary Figure S1.** Top panel: sequence similarity network of *Streptococcus* RiPP BGCs based on TLAT sequences. Nodes represent gene clusters and are colored/grouped by similarity ((edge % identity of 20). Bottom panel: representative gene clusters from each of the 11 groupings. NCBI accession codes are included for all putative enzymes.

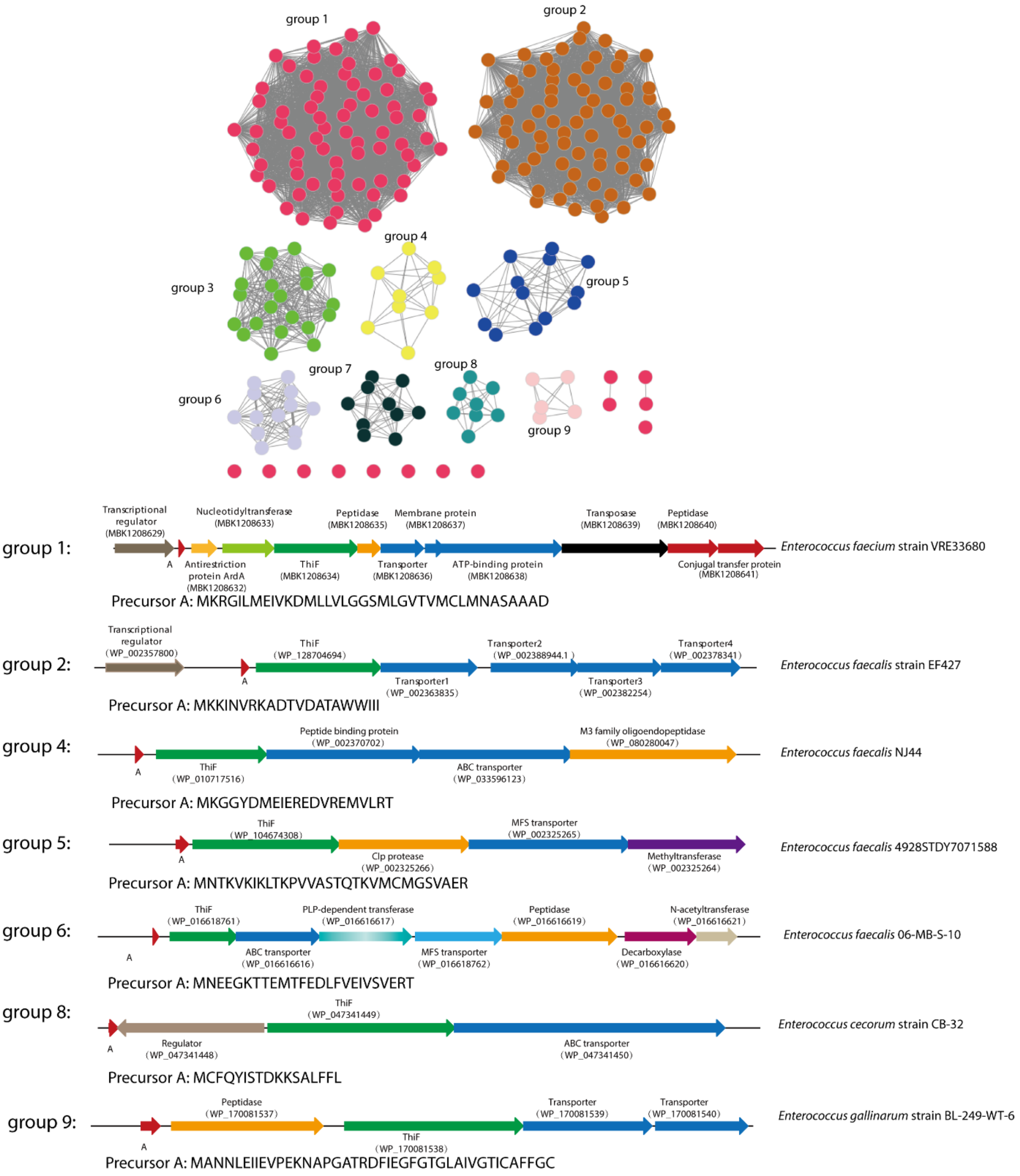


**Supplementary Figure S2.** Top panel: sequence similarity network of *Enterococcus* RiPP BGCs based on TLAT sequences. Nodes represent gene clusters and are colored/grouped by similarity((edge % identity of 20). Bottom panel: representative gene clusters from each of the 9 groupings. NCBI accession codes are included for all putative enzymes.

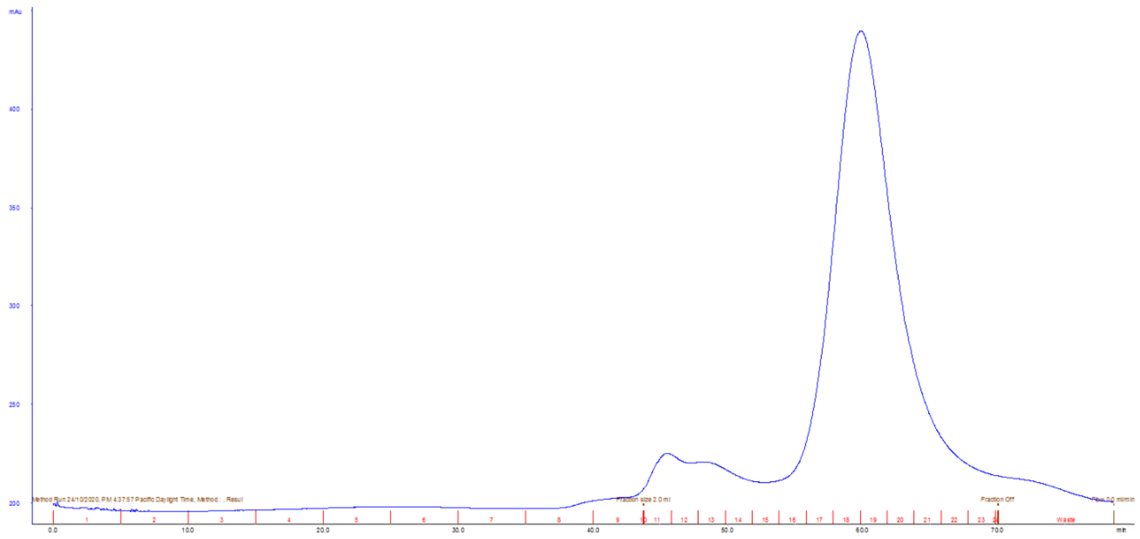

**Supplementary Figure S3.** Purification of EnfB by size-exclusion chromatography.Fractions(18-20) were combined and further collected.

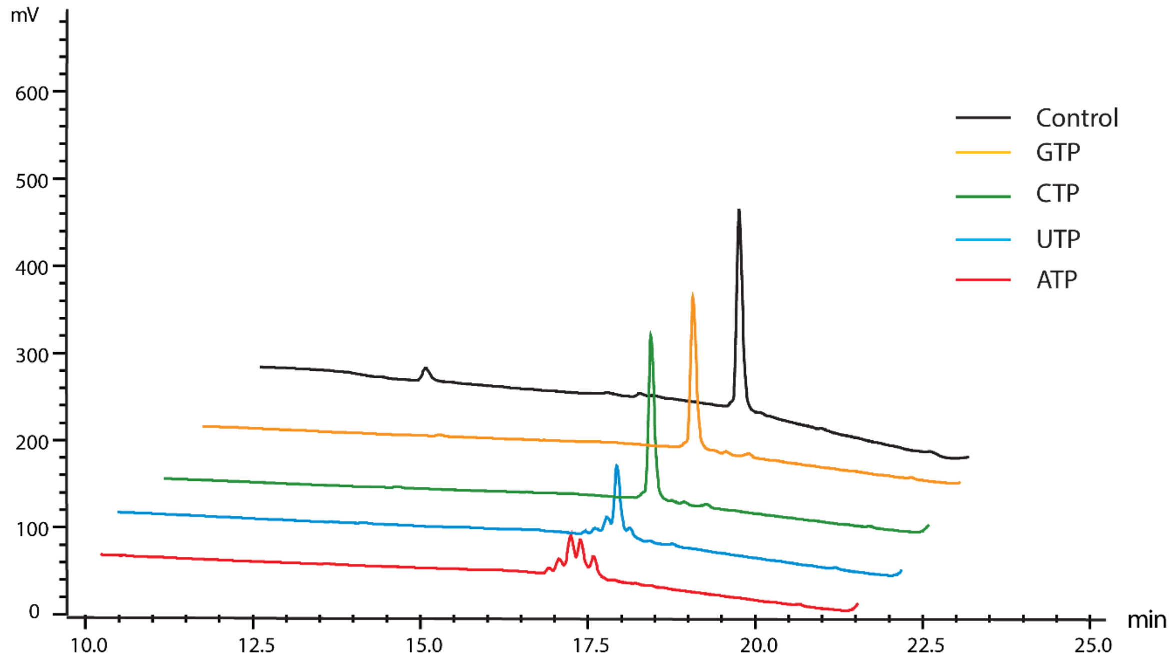


**Supplementary Figure S4.** Stacked and staggered HPLC chromatograms of *in vitro* assays of EnfB with EnfA and GTP, CTP, UTP, or ATP.

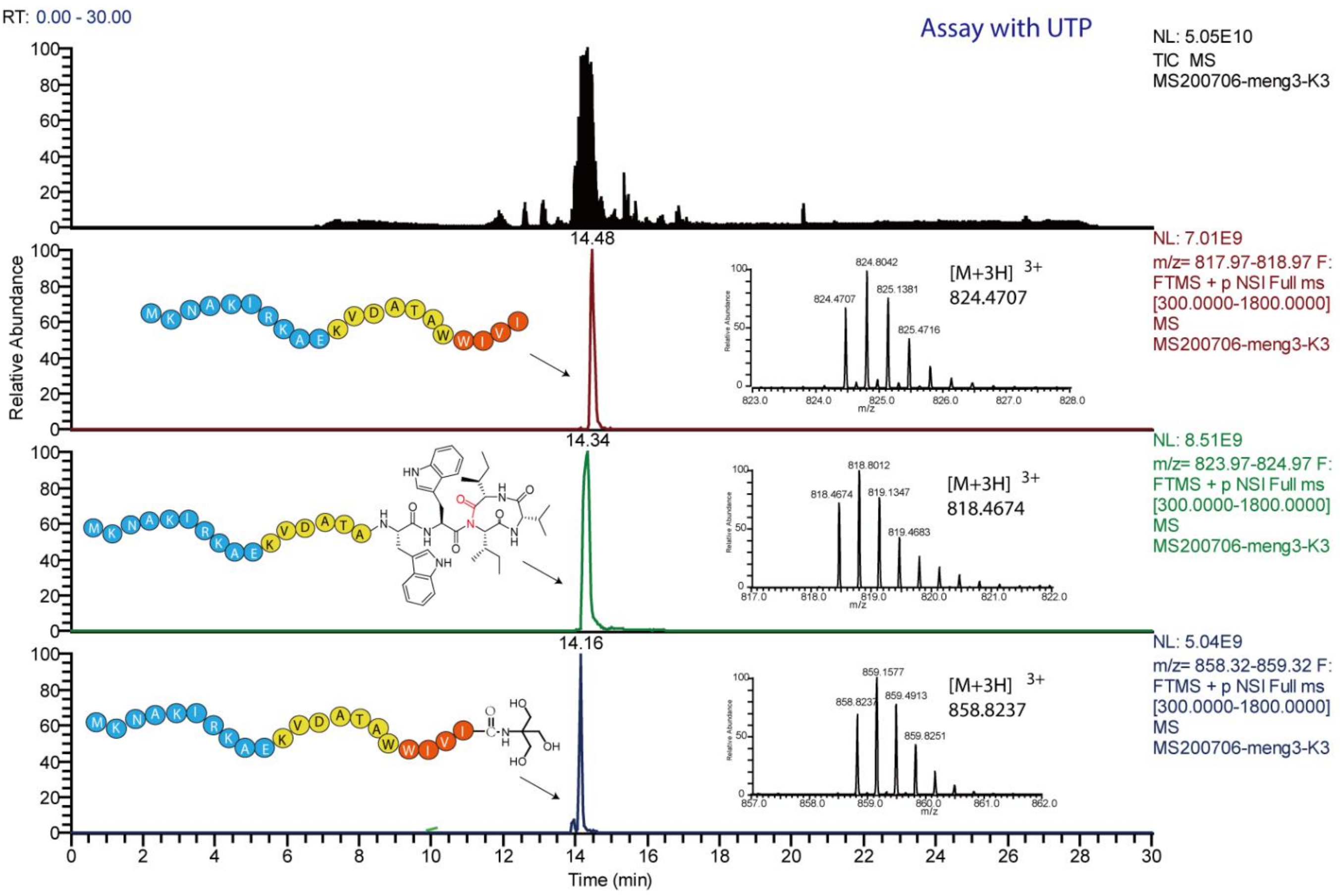


**Supplementary Figure S5.** Extracted ion chromatograms and mass spectra from the assay of EnfB with EnfA and UTP in Tris buffer.

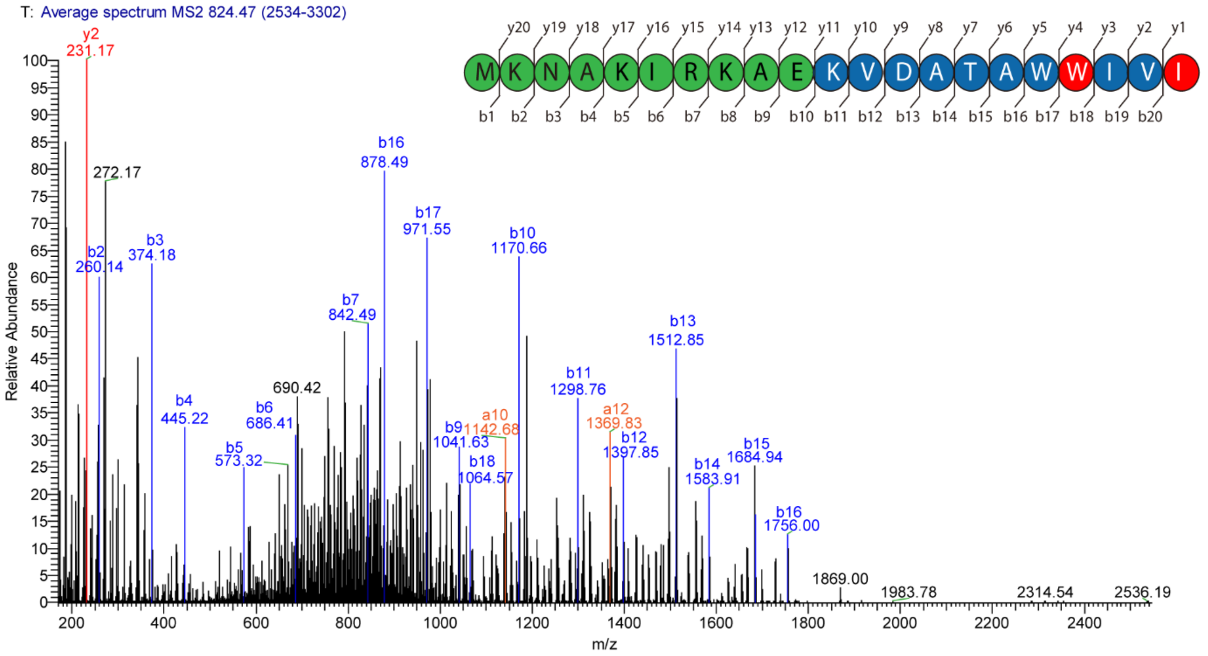
T: Average spectrum MS2 824.47 (2534-3302)
y20 y19 y18 y17 y16 y15 y14 y13 y12 y11 y10 y9 y8 y7 y6 y5 y4 y3 y2 y1
M K N A K I R K A E K V D A T A W W I V I
b1 b2 b3 b4 b5 b6 b7 b8 b9 b10 b11 b12 b13 b14 b15 b16 b17 b18 b19 b20
y2 231.17
272.17
b2 260.14
b3 374.18
b4 445.22
b5 573.32
b6 686.41
690.42
b7 842.49
b16 878.49
b17 971.55
b9 1041.63
b18 1064.57
a10 1142.68
b10 1170.66
b11 1298.76
a12 1369.83
b12 1397.85
b13 1512.85
b14 1583.91
b15 1684.94
b16 1756.00
1869.00
1983.78
2314.54
2536.19
Relative Abundance
m/z

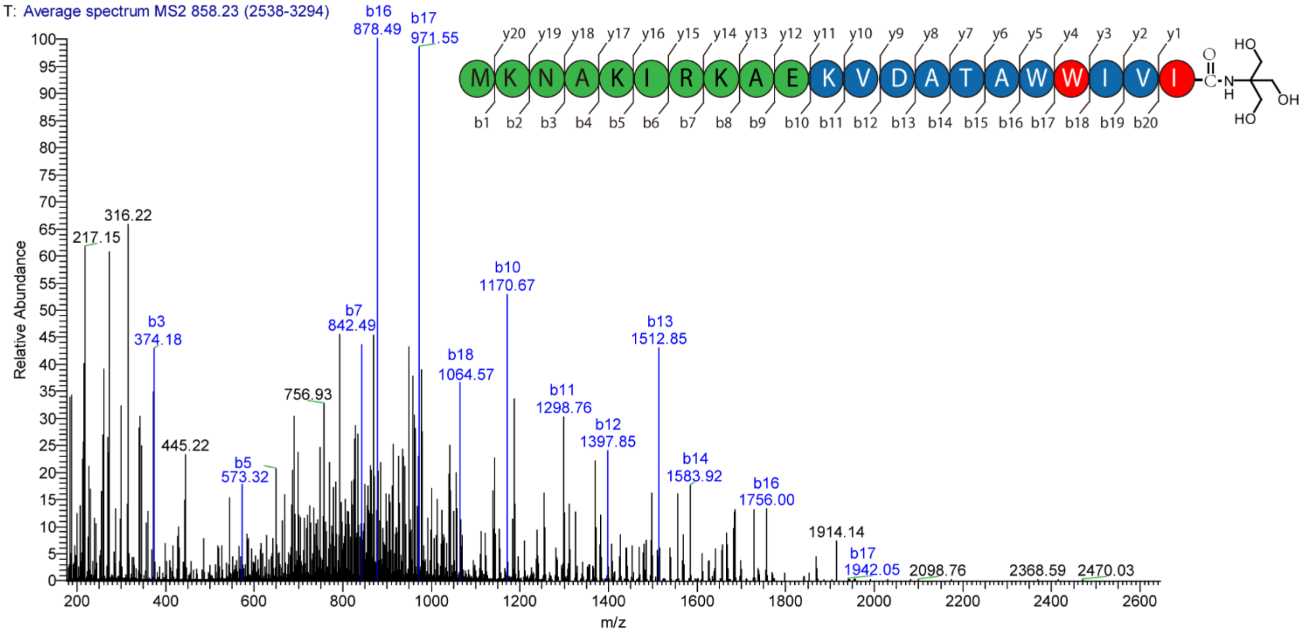
T: Average spectrum MS2 858.23 (2538-3294)
y20 y19 y18 y17 y16 y15 y14 y13 y12 y11 y10 y9 y8 y7 y6 y5 y4 y3 y2 y1
M K N A K I R K A E K V D A T A W W I V I
b1 b2 b3 b4 b5 b6 b7 b8 b9 b10 b11 b12 b13 b14 b15 b16 b17 b18 b19 b20
HO
OH
HO
217.15
316.22
b3 374.18
445.22
b5 573.32
756.93
b7 842.49
b16 878.49
b17 971.55
b18 1064.57
b10 1170.67
b11 1298.76
b12 1397.85
b13 1512.85
b14 1583.92
b16 1756.00
1914.14
b17 1942.05
2098.76
2368.59
2470.03
Relative Abundance
m/z

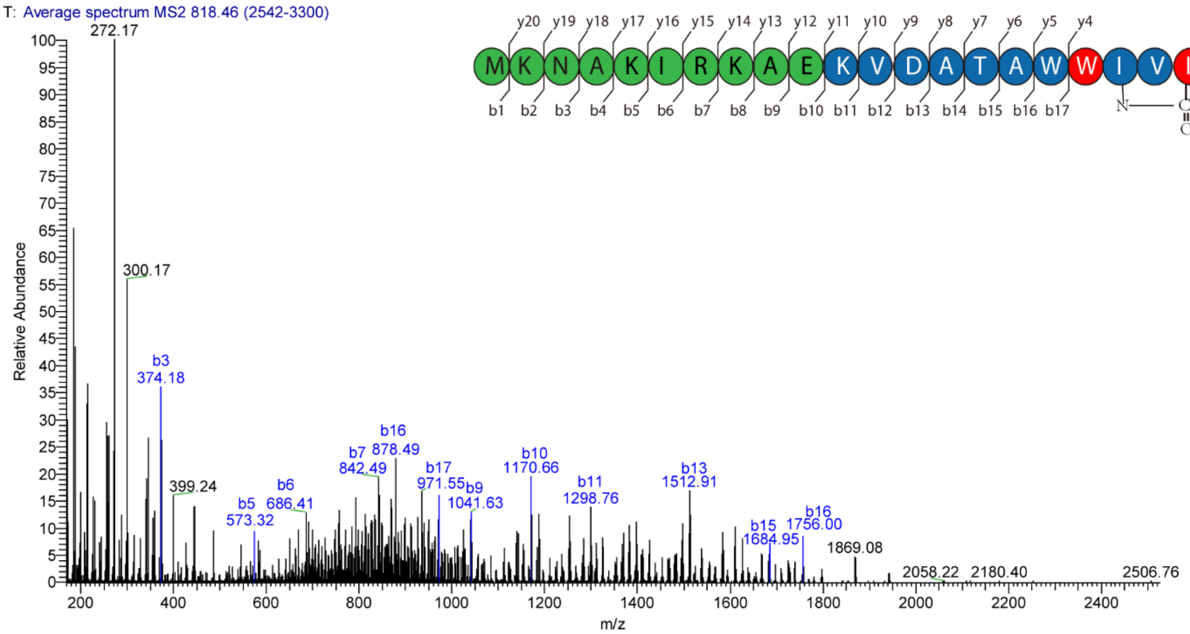
T: Average spectrum MS2 818.46 (2542-3300)
y20 y19 y18 y17 y16 y15 y14 y13 y12 y11 y10 y9 y8 y7 y6 y5 y4
M K N A K I R K A E K V D A T A W W I V I
b1 b2 b3 b4 b5 b6 b7 b8 b9 b10 b11 b12 b13 b14 b15 b16 b17
272.17
300.17
b3 374.18
399.24
b5 573.32
b6 686.41
b7 842.49
b16 878.49
b17 971.55
b9 1041.63
b10 1170.66
b11 1298.76
b13 1512.91
b15 1684.95
b16 1756.00
1869.08
2058.22
2180.40
2506.76
Relative Abundance
m/z

**Supplementary Figure S6.** MS/MS fragmentation of peptides in the enzymatic assays. Fragments of the b-series are highlighted in blue, with those the y-series in red.

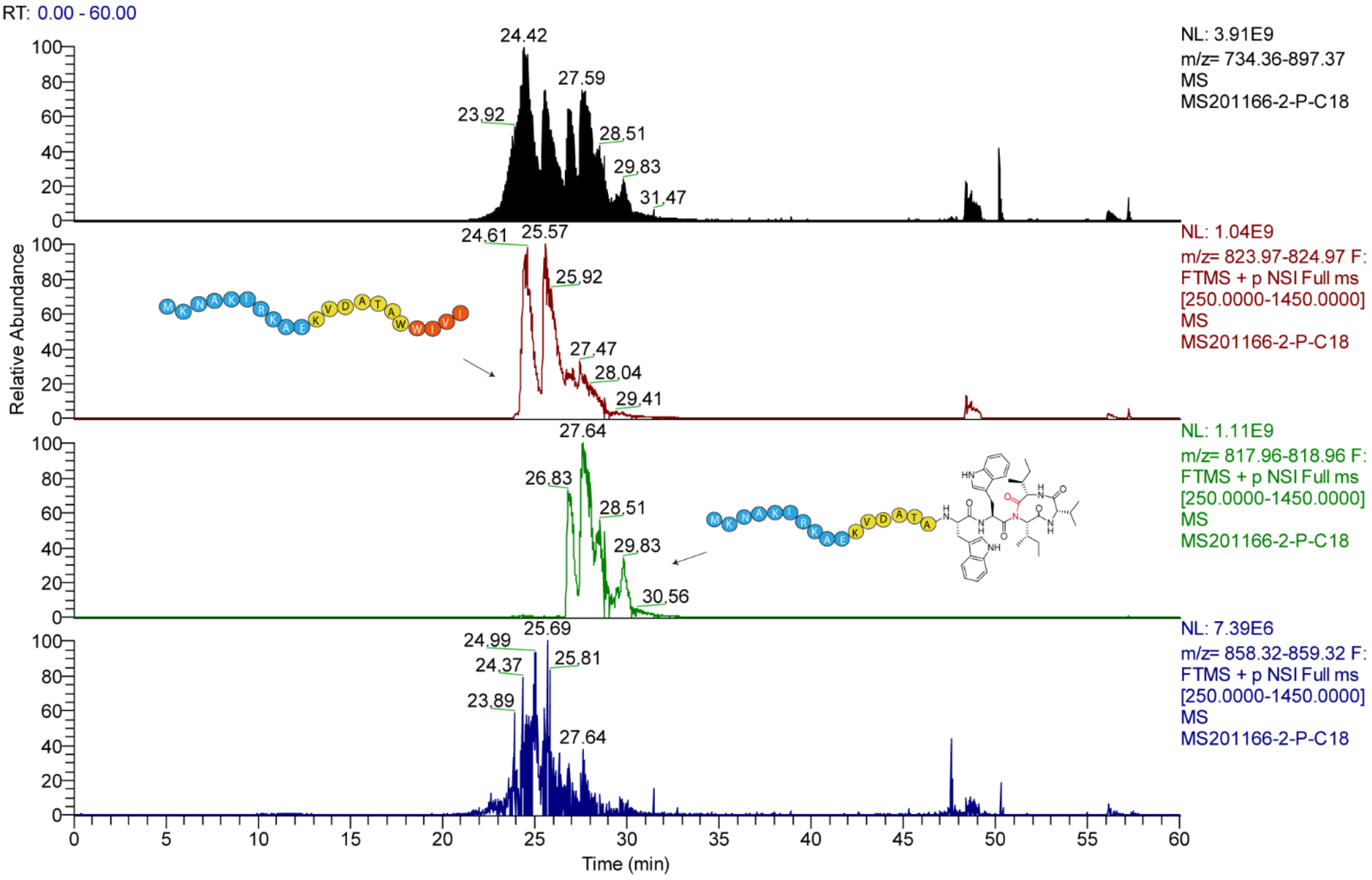


**Supplementary Figure S7.** Extracted ion chromatograms and mass spectra from the assay of EnfB with EnfA and ATP in Tris buffer.

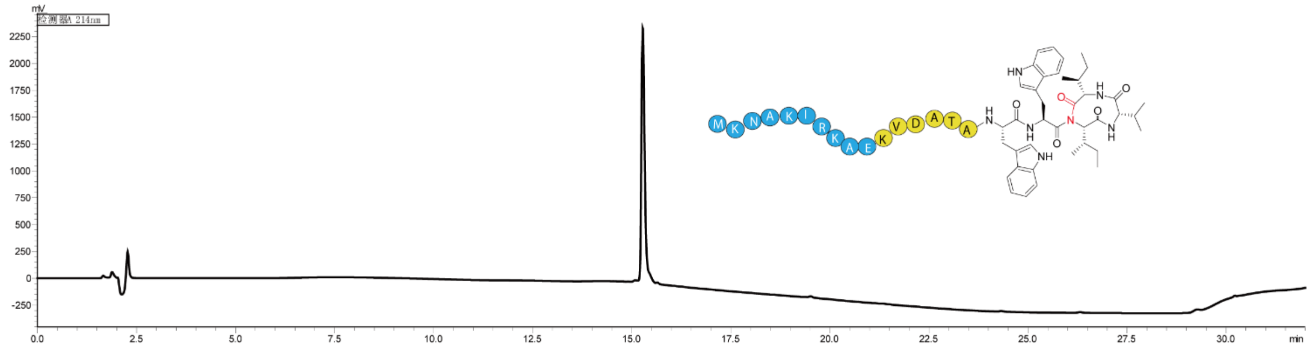

**Supplementary Figure S8.** Purification of enterofaecin by preparative HPLC.

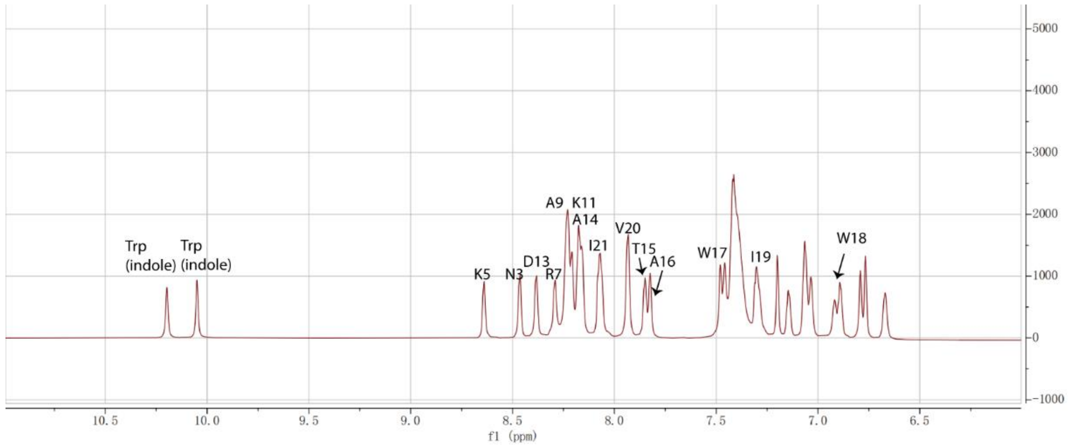


**Supplementary Figure S9.** [1]H NMR spectrum of linear EnfA in the range of 11.0 to 6.0 ppm.

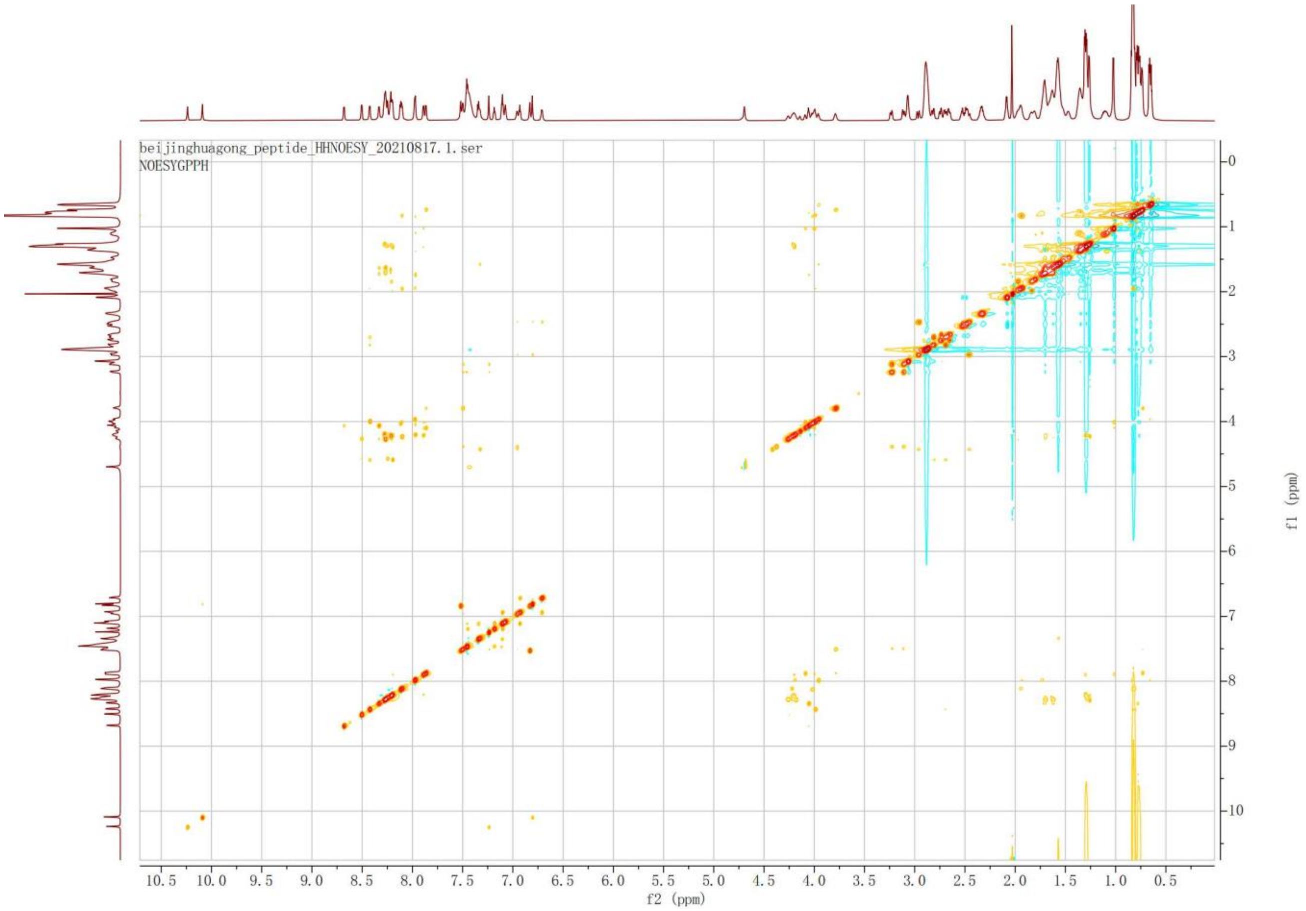


**Supplementary Figure S10.** Section of NOESY spectrum of linear EnfA.

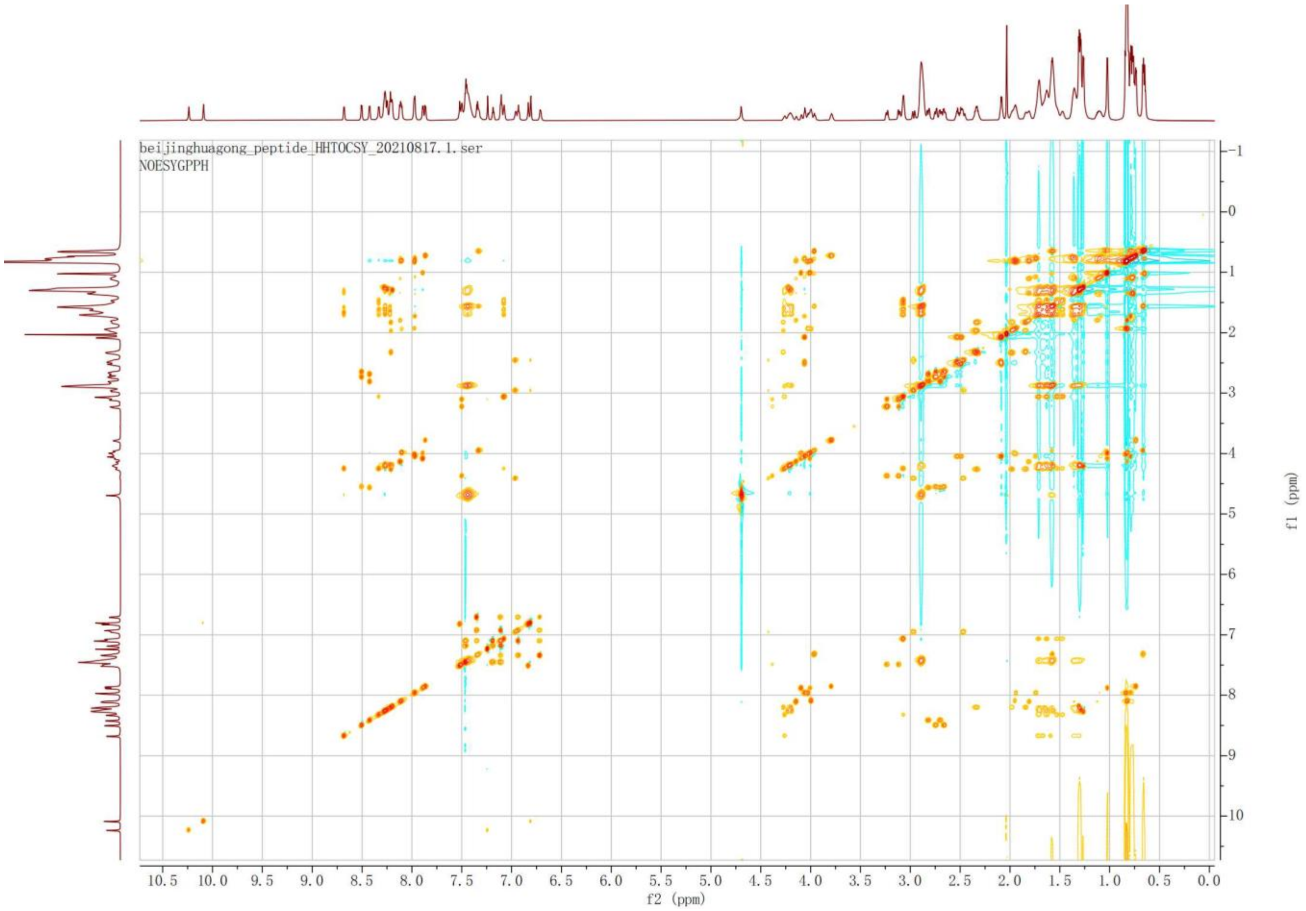


**Supplementary Figure S11.** Section of TOCSY spectrum of linear EnfA

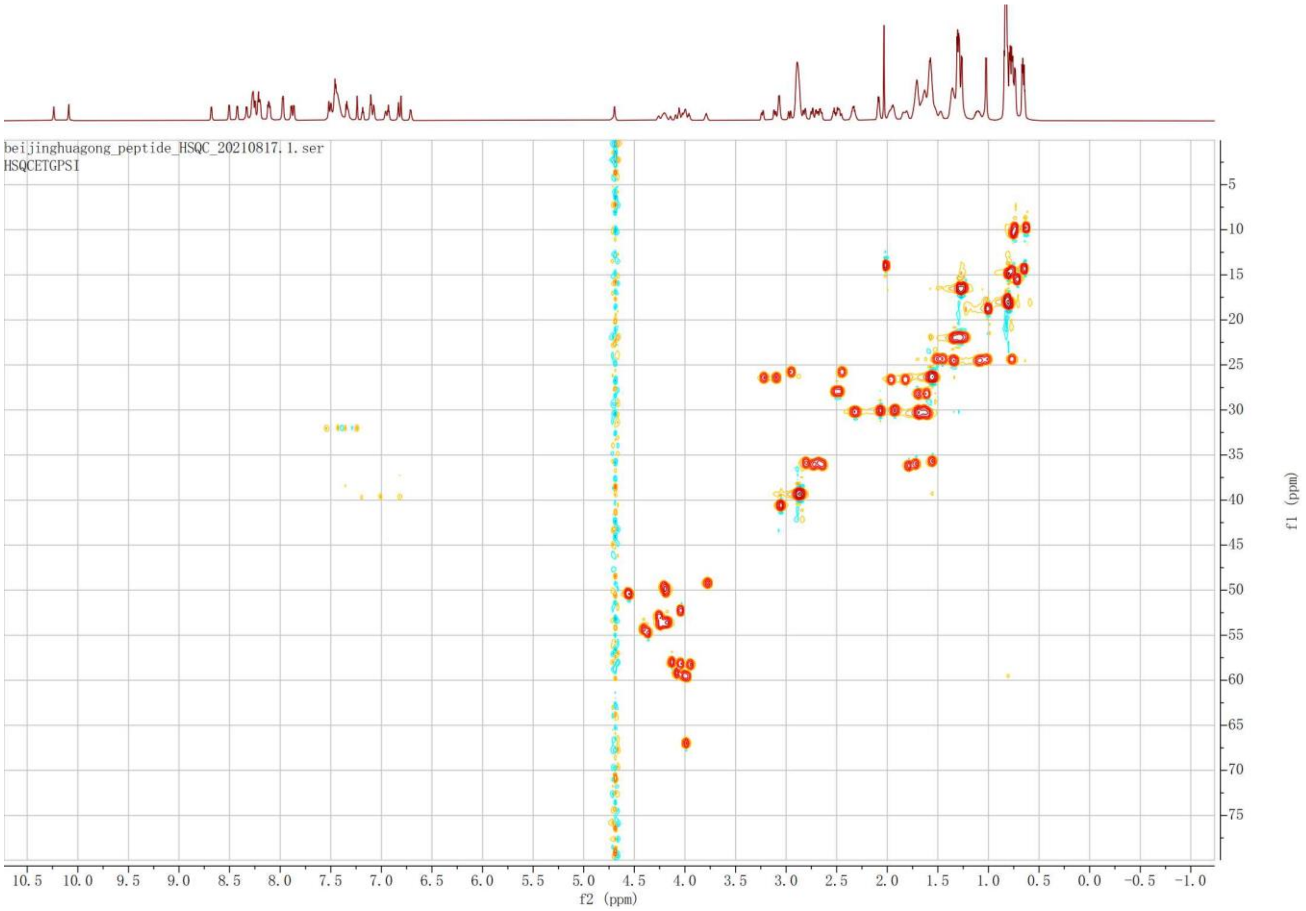


**Supplementary Figure S12.** Section of HSQC spectrum of linear EnfA.

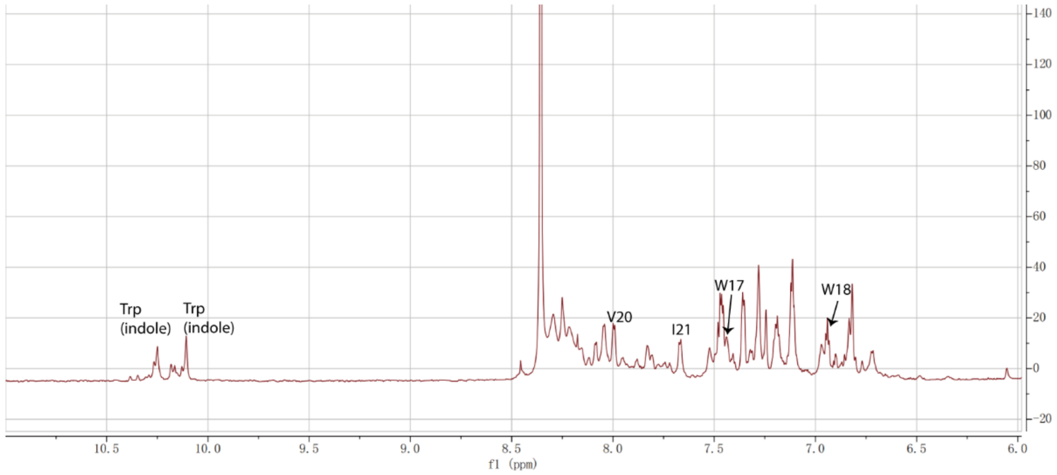


**Supplementary Figure S13.** $^1$H NMR spectrum of enterofaecin in the range of 11.0 to 6.0 ppm

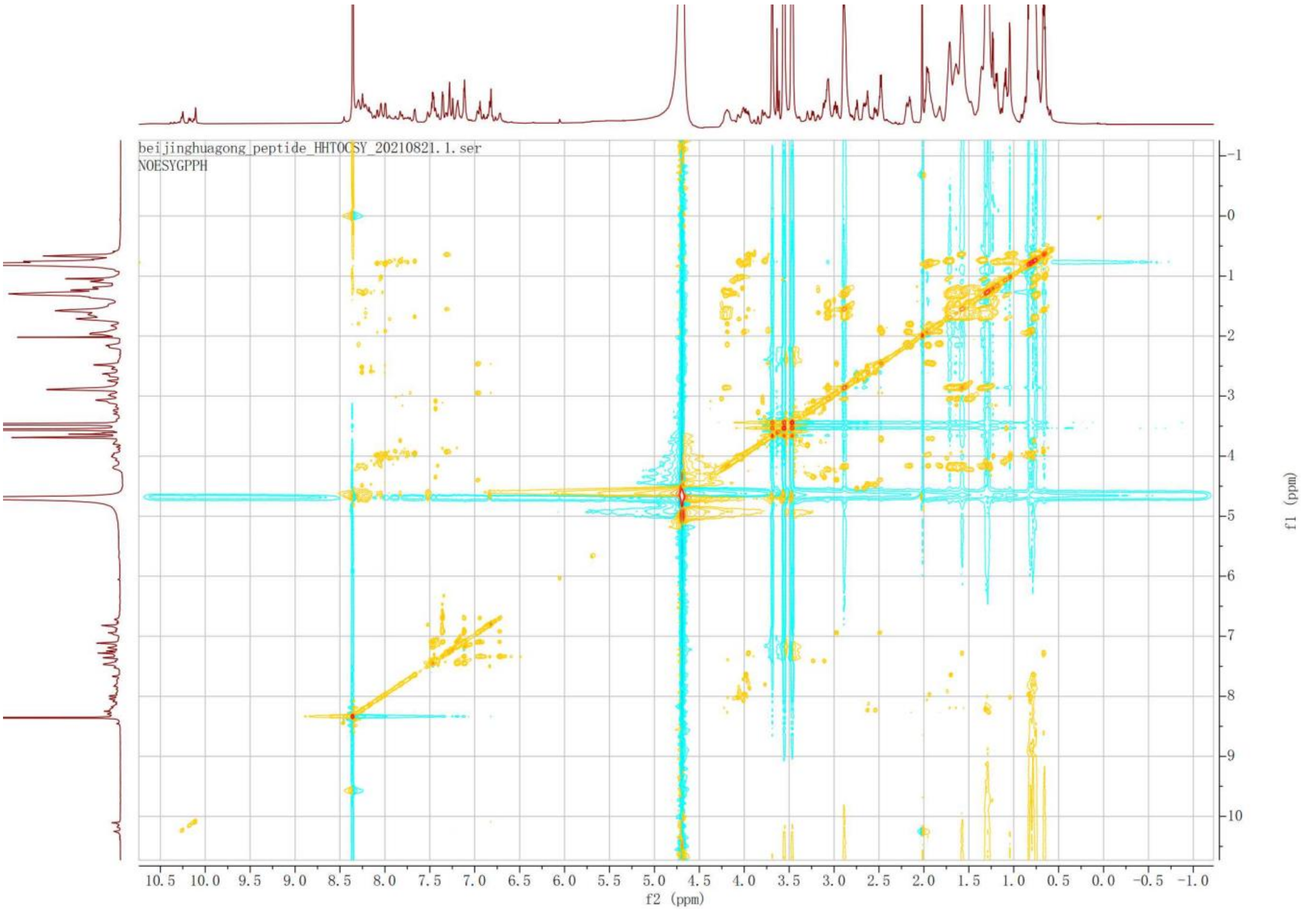


**Supplementary Figure S14.** Section of TOCSY spectrum of enterofaecin.

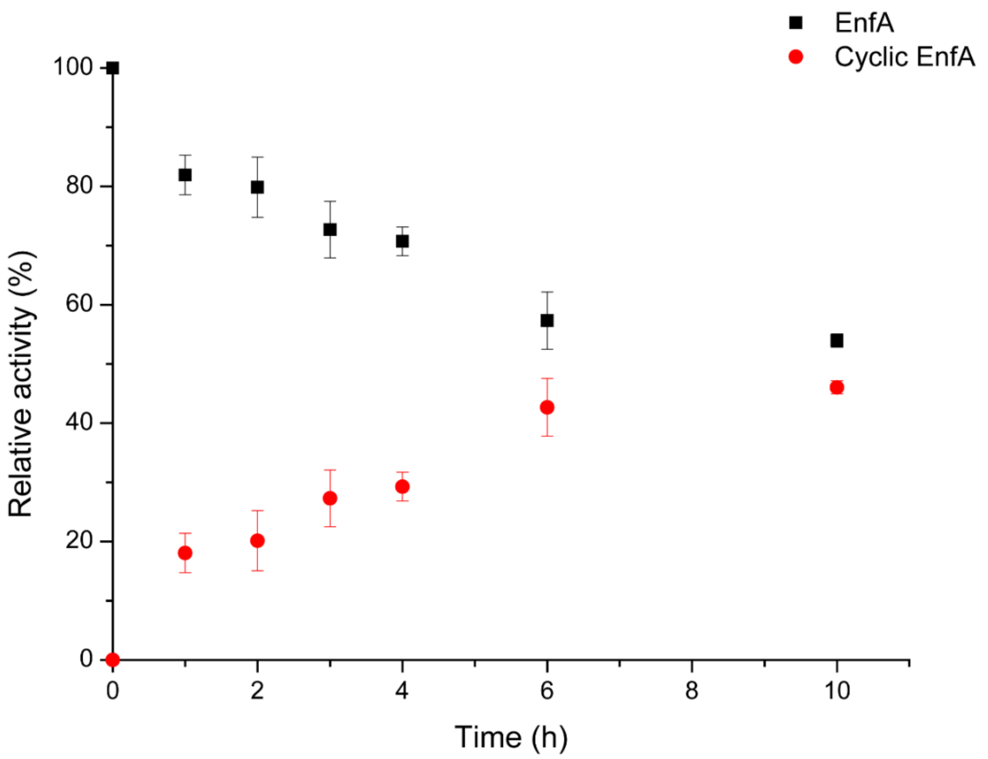


**Supplementary Figure S15.** Time-dependent conversion of EnfA to enterofaecin (cyclic EnfA) as catalyzed by EnfB.Error bars are from s.d. from triplicate measurements.

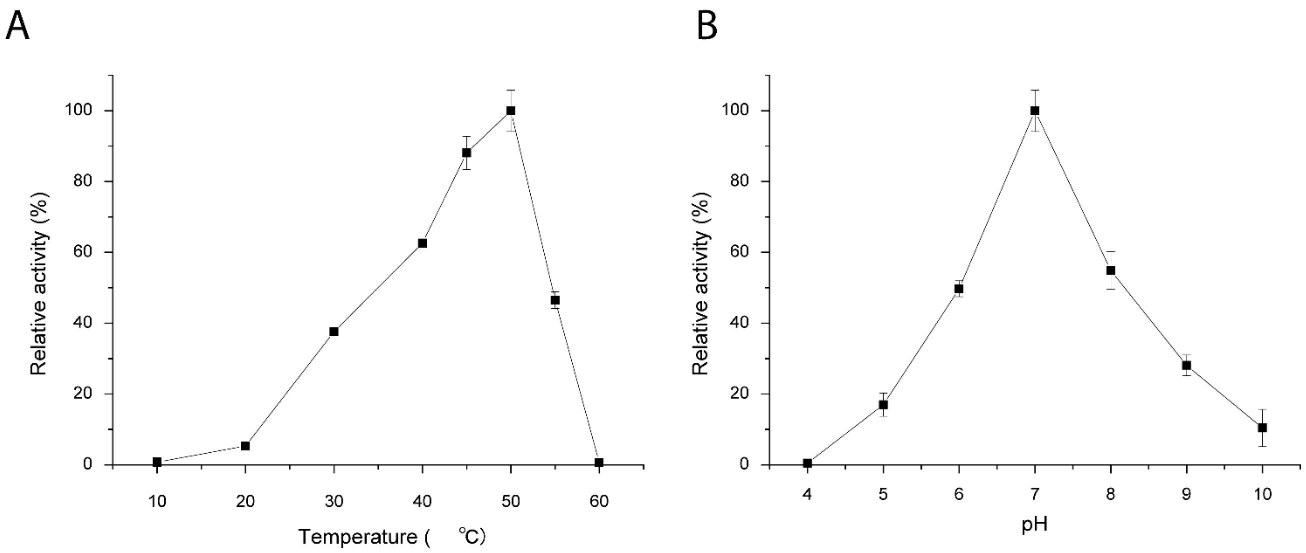


**Supplementary Figure S16.** (A). Effects of the temperature on EnfB activity. The activity at 50 °C was taken as 100%. (B). Effects of pH on EnfB activity. The activity at pH 7.0 was taken as 100%. Each point represents the mean ± s.d. from three independent experiments.

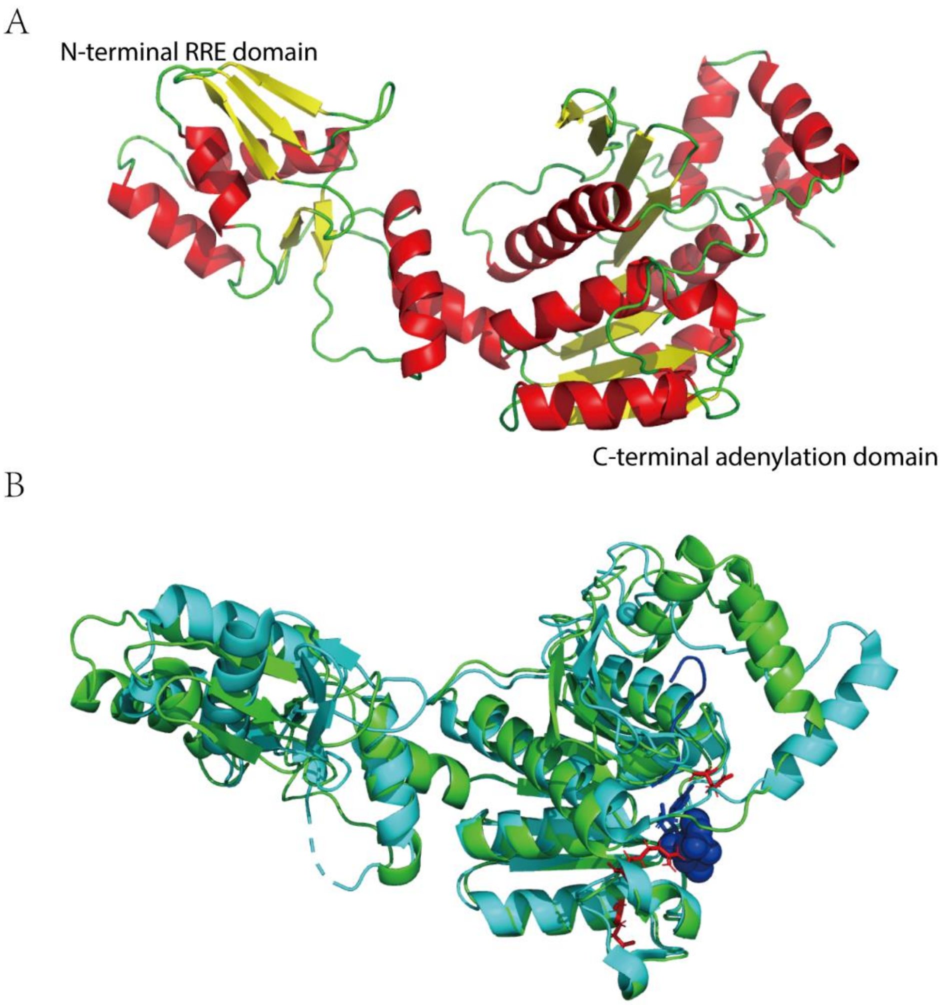


**Supplementary Figure S17.** (A) AlphaFold homology model of EnfB. The N-terminal RRE domain and C-terminal adenylation domain are labeled. (B) AlphaFold homology model of EnfB (green) overlaid with the structure of MccB (PDB ID 6OM4), which participates in the biosynthesis of the antibiotic microcin C7.

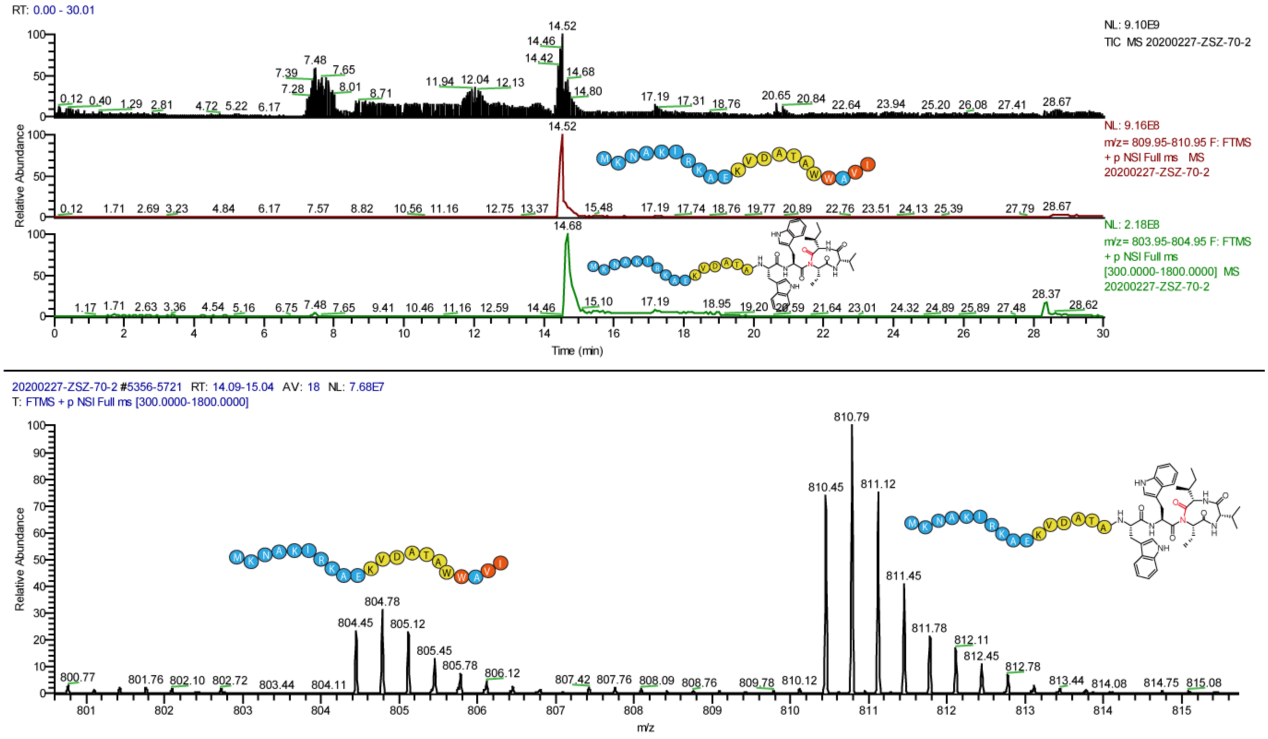


**Supplementary Figure S18.** MS analysis of *in vitro* assays with EnfB and EnfA$_{I19A}$.

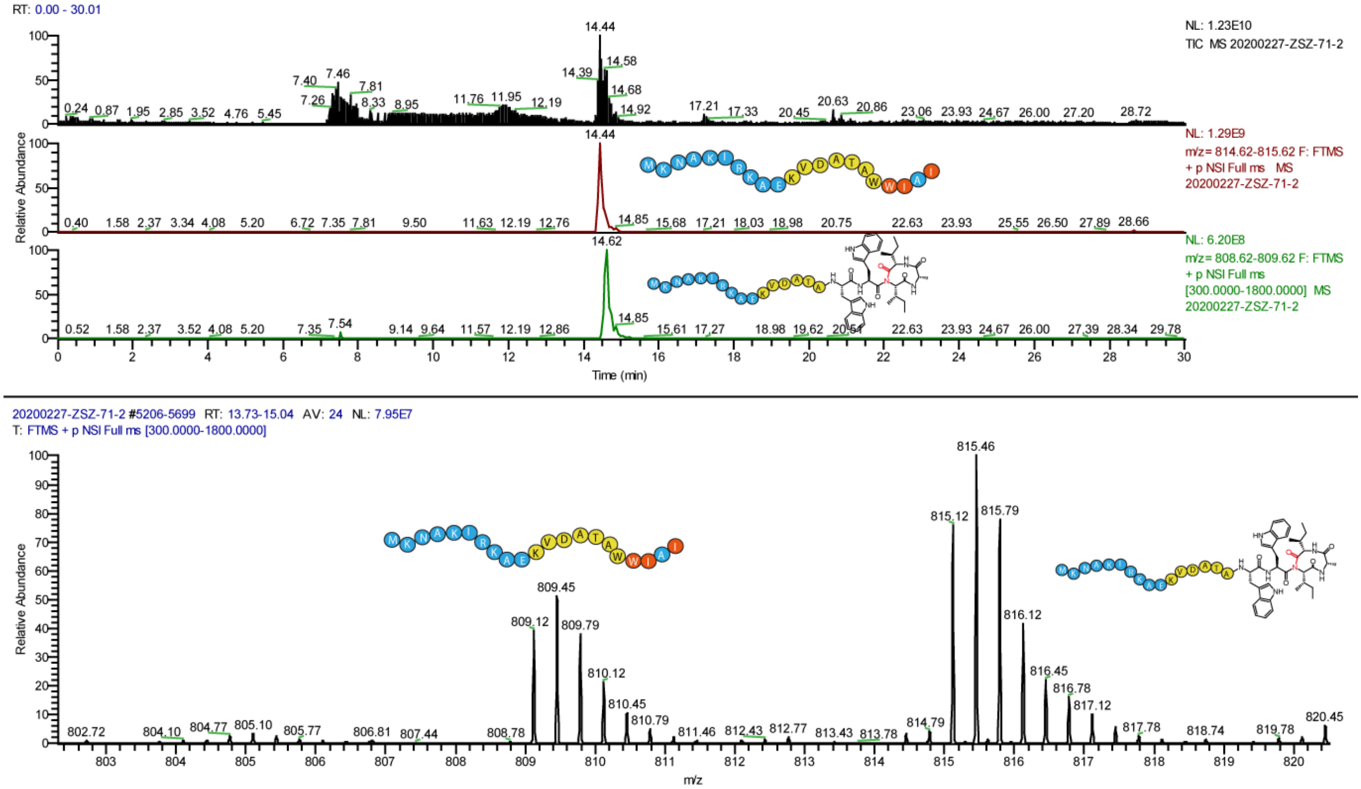


**Supplementary Figure S19.** MS analysis of *in vitro* assays with EnfB and EnfA$_{V20A}$.

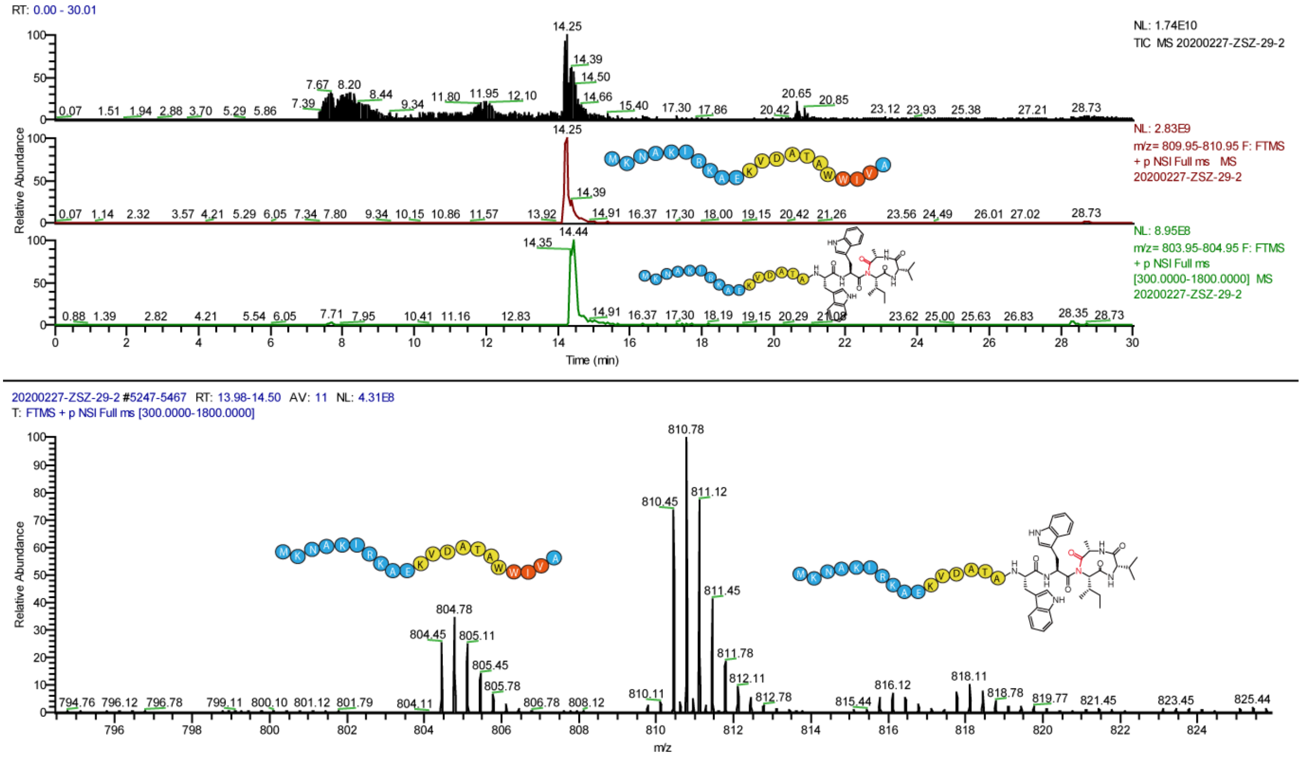

**Supplementary Figure S20.** MS analysis of *in vitro* assays with EnfB and EnfA$_{I21A}$.

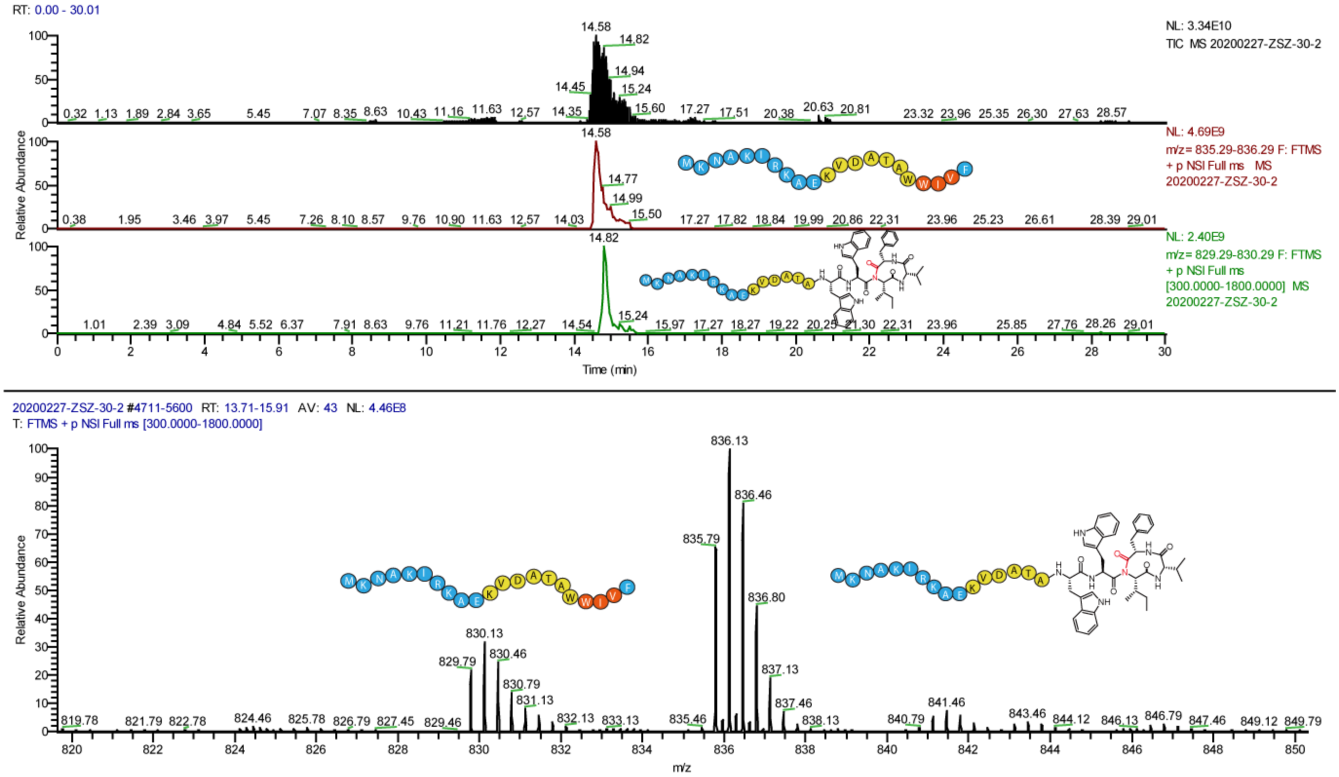

**Supplementary Figure S21.** MS analysis of *in vitro* assays with EnfB and EnfA$_{I21F}$.

A

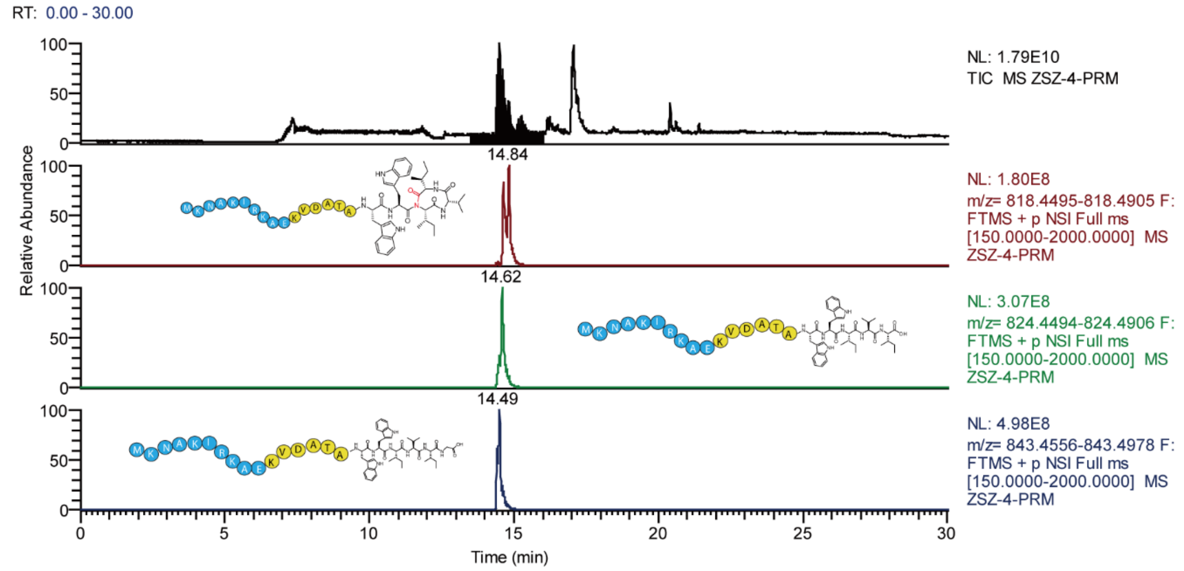


B

### Mass/Charge Table

| | Mass | |
|---|---|---|
| | Mono | Avg |
| (M) | 2527.40829 | 2529.00972 |
| $(M+H)^{+}$ | 2528.41557 | 2530.01700 |
| $(M+2H)^{2+}$ | 1264.71145 | 1265.51216 |
| $(M+3H)^{3+}$ | 843.47674 | 844.01055 |
| $(M+4H)^{4+}$ | 632.85938 | 633.25974 |

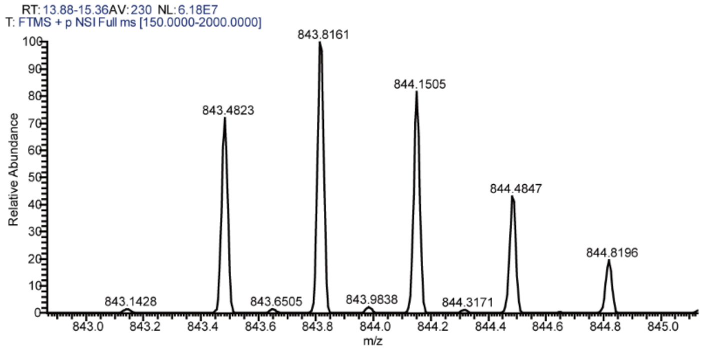


* C-terminus modification: 57.02

C

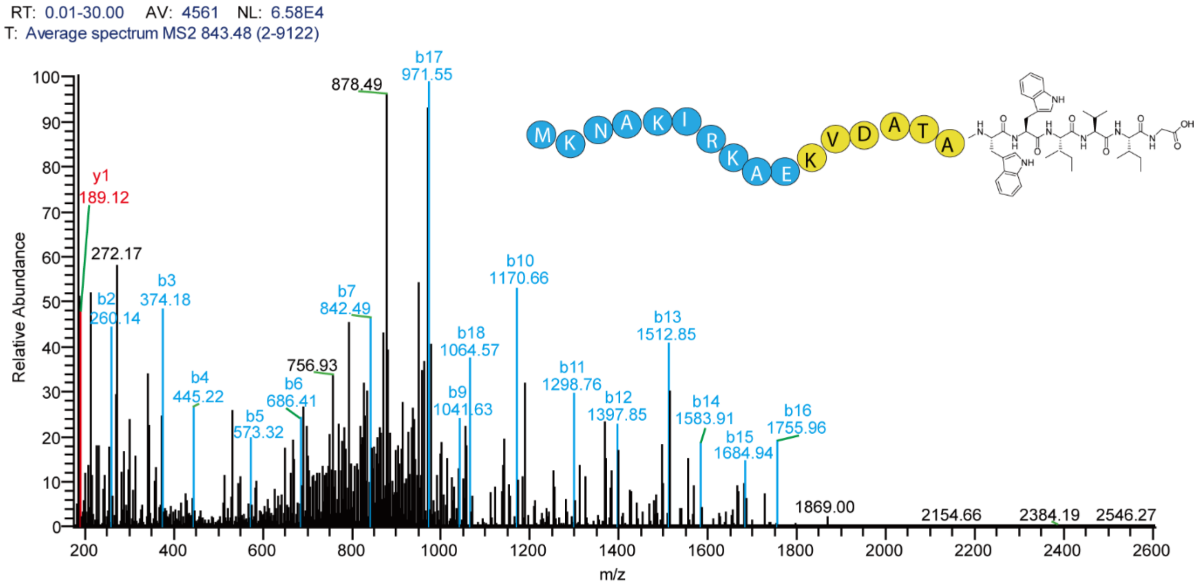


**Supplementary Figure S22.** MS/MS fragmentation of peptides in the enzymatic assays with glycine. Fragments of the b-series are highlighted in blue, with those of the y-series in red.

A

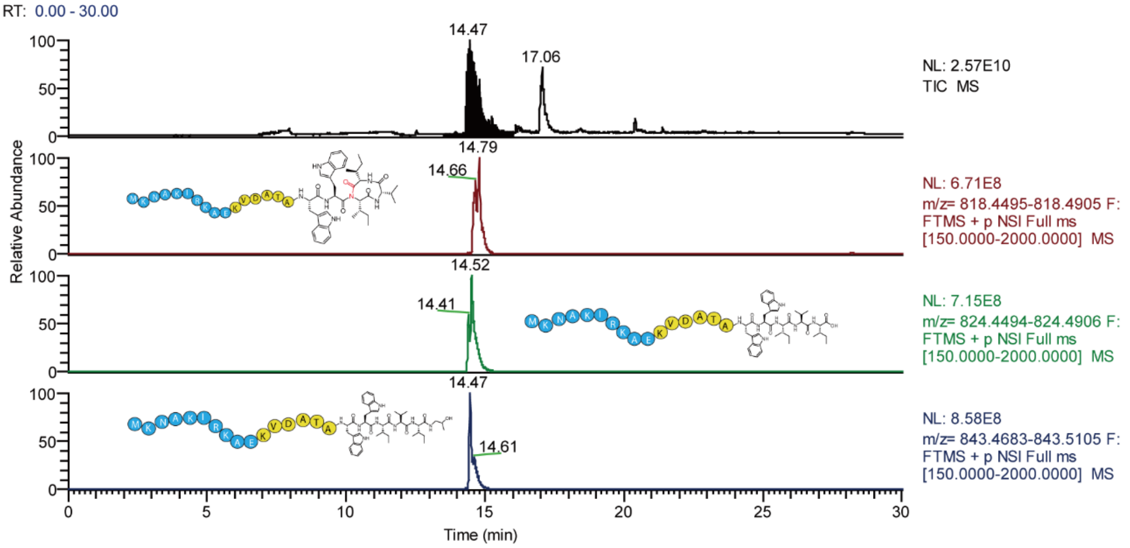


B

**Mass/Charge Table**

| | Mass | |
|---|---|---|
| | Mono | Avg |
| (M) | 2527. 44629 | 2529. 04773 |
| (M+H)$^{+}$ | 2528. 45357 | 2530. 05500 |
| (M+2H)$^{2+}$ | 1264. 73045 | 1265. 53116 |
| (M+3H)$^{3+}$ | 843. 48940 | 844. 02321 |
| (M+4H)$^{4+}$ | 632. 86888 | 633. 26924 |

* C-terminus modification: 57.058

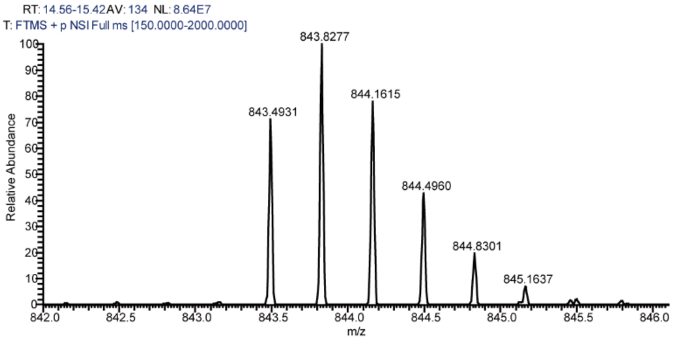


C

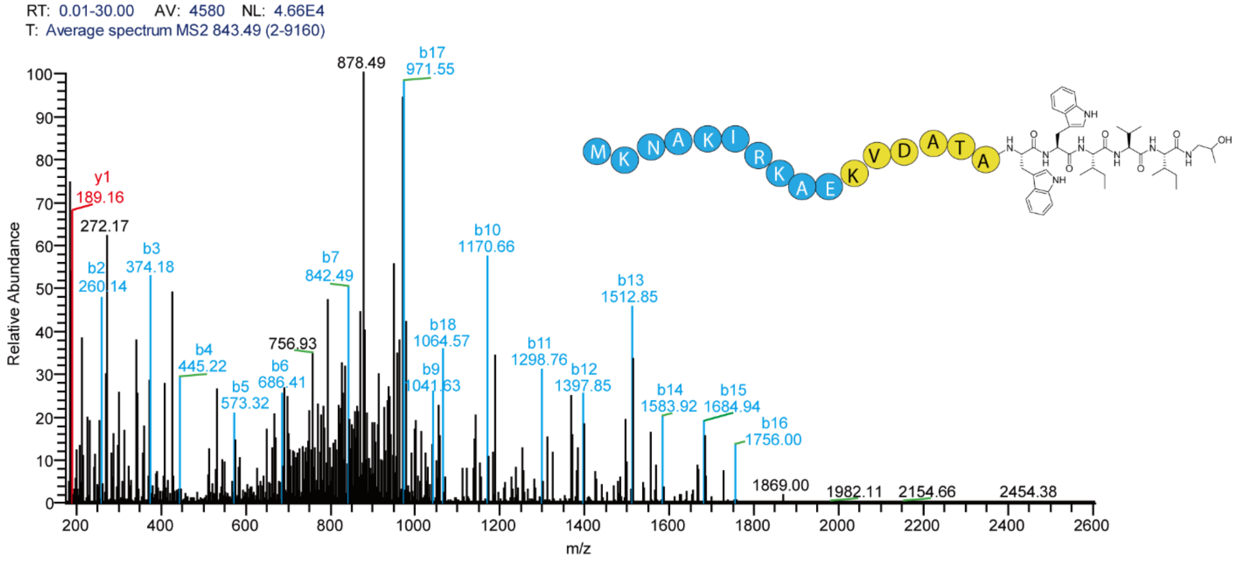


**Supplementary Figure S23.** MS/MS fragmentation of peptides in the enzymatic assays with amino-2-propanol. Fragments of the b-series are highlighted in blue, with those of the y-series in red.

A

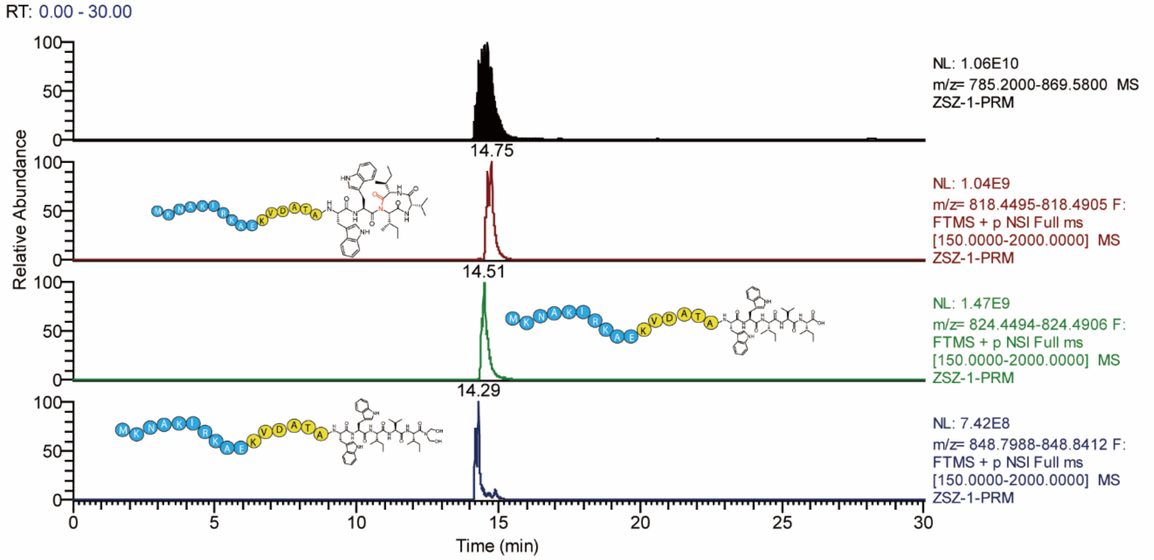


B

**Mass/Charge Table**

| | Mass | |
|---|---|---|
| | Mono | Avg |
| (M) | 2543. 43829 | 2545. 03973 |
| (M+H)$^{+}$ | 2544. 44557 | 2546. 04700 |
| (M+2H)$^{2+}$ | 1272. 72645 | 1273. 52716 |
| (M+3H)$^{3+}$ | 848. 82007 | 849. 35388 |
| (M+4H)$^{4+}$ | 636. 86688 | 637. 26724 |

* C-terminus modification: 73.05

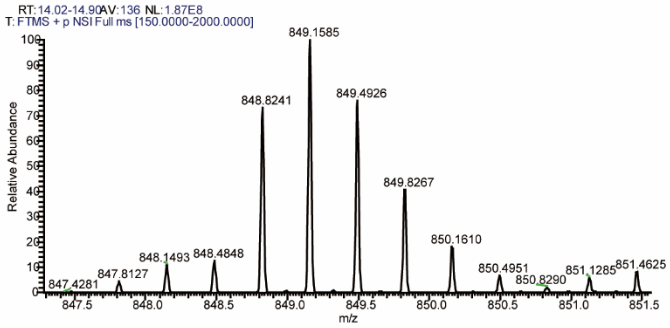


C

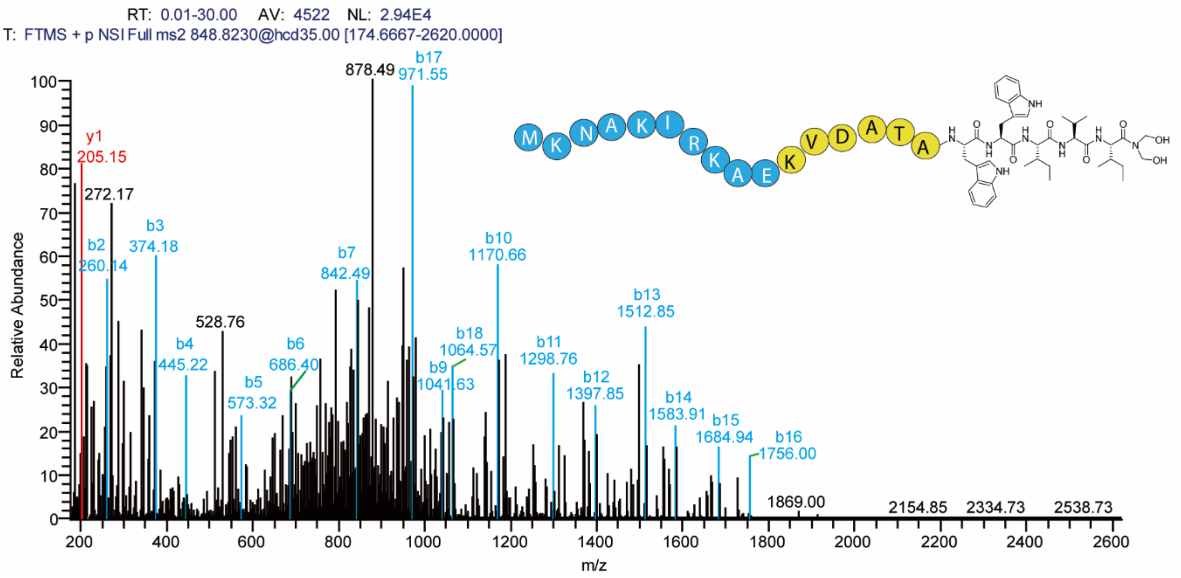


**Supplementary Figure S24.** MS/MS fragmentation of peptides in the enzymatic assays with 2-amino-1,3-propanediol. Fragments of the b-series are highlighted in blue, with those of the y-series in red.

A

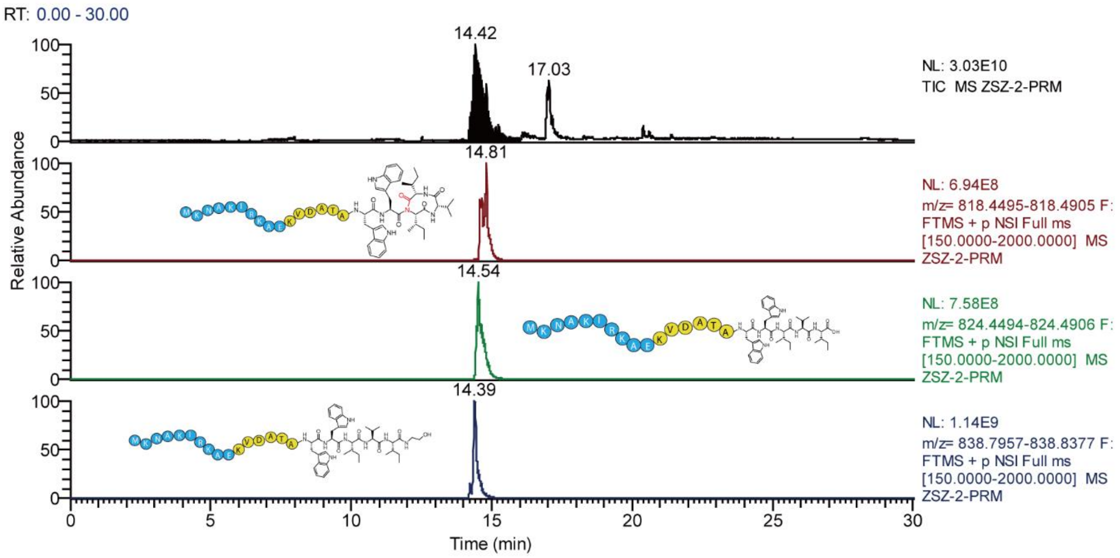


B

**Mass/Charge Table**

| | Mass | |
|---|---|---|
| | **Mono** | **Avg** |
| **(M)** | 2513. 42829 | 2515. 02972 |
| **(M+H)$^{+}$** | 2514. 43557 | 2516. 03700 |
| **(M+2H)$^{2+}$** | 1257. 72145 | 1258. 52216 |
| **(M+3H)$^{3+}$** | 838. 81674 | 839. 35055 |
| **(M+4H)$^{4+}$** | 629. 36438 | 629. 76474 |

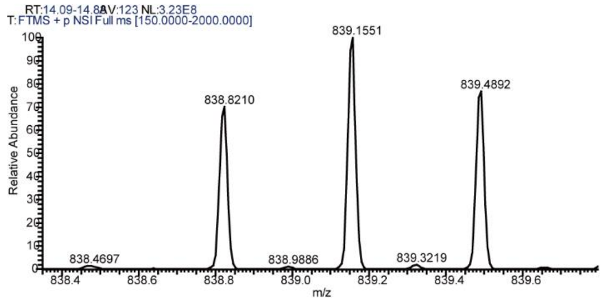


* C-terminus modification: 43.04

C

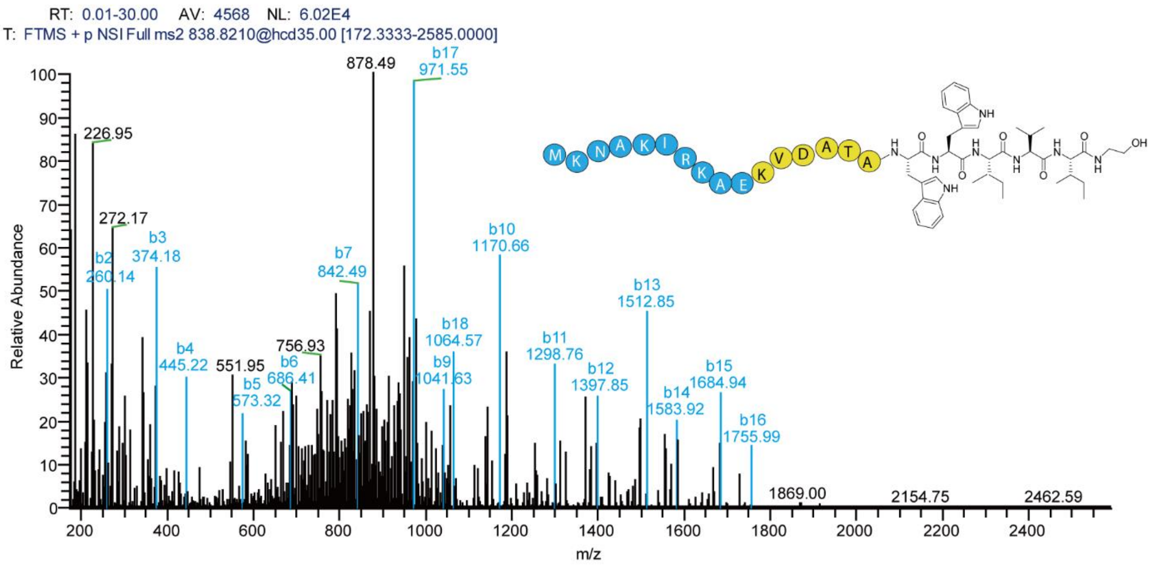


**Supplementary Figure S25.** MS/MS fragmentation of peptides in the enzymatic assays with ethanolamine. Fragments of the b-series are highlighted in blue, with those of the y-series in red.

A

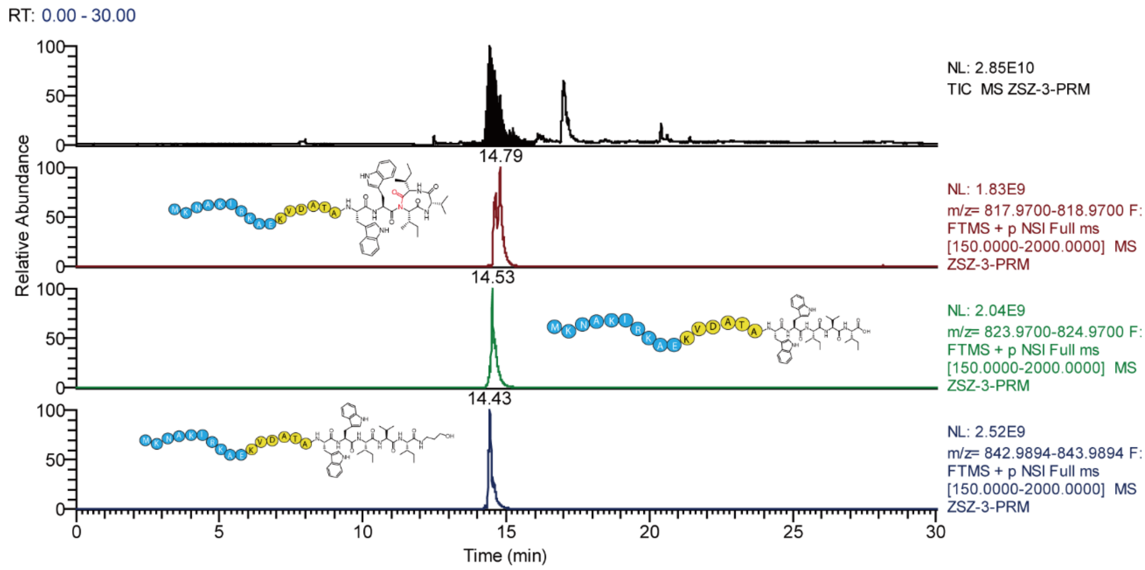


B

**Mass/Charge Table**

| | Mass | |
|---|---|---|
| | **Mono** | **Avg** |
| **(M)** | 2527. 44629 | 2529. 04773 |
| **(M+H)⁺** | 2528. 45357 | 2530. 05500 |
| **(M+2H)²⁺** | 1264. 73045 | 1265. 53116 |
| **(M+3H)³⁺** | 843. 48940 | 844. 02321 |
| **(M+4H)⁴⁺** | 632. 86888 | 633. 26924 |

* C-terminus modification: 57.058

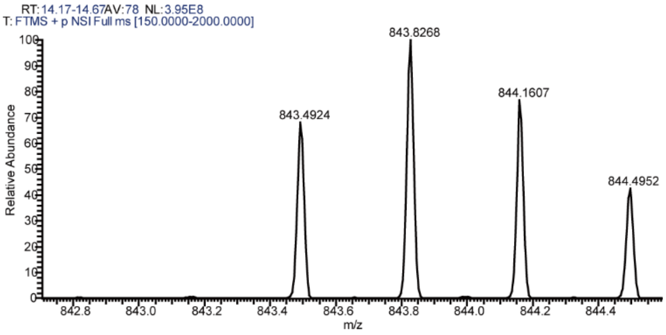


C

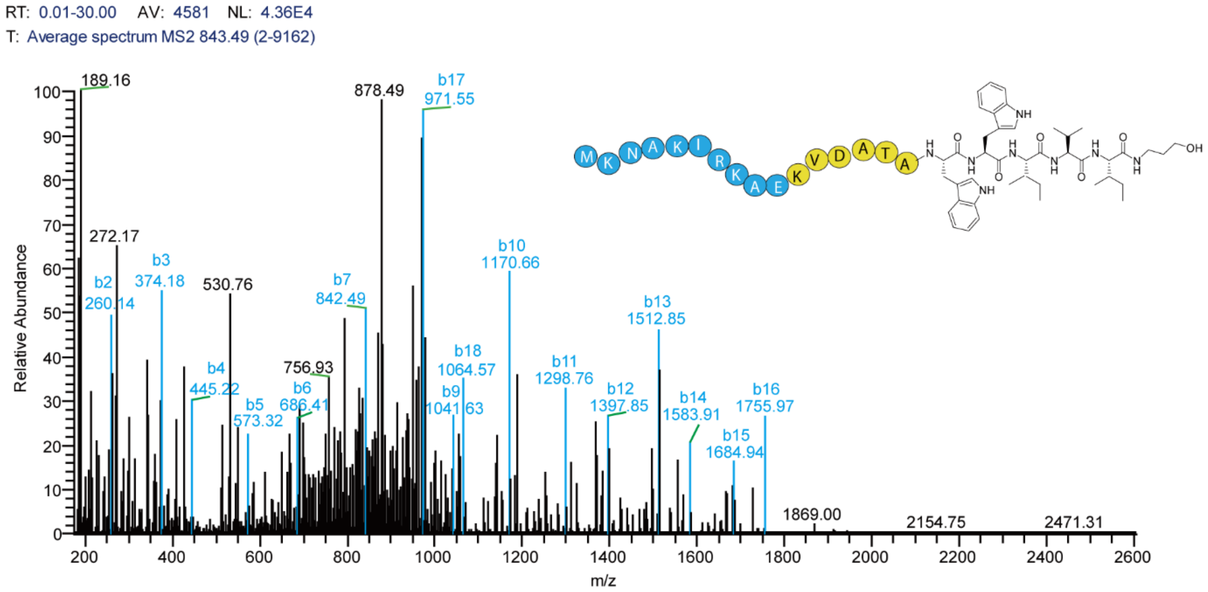


**Supplementary Figure S26.** MS/MS fragmentation of peptides in the enzymatic assays with 3-amino-1-propanol. Fragments of the b-series are highlighted in blue, with those of the y-series in red.

A

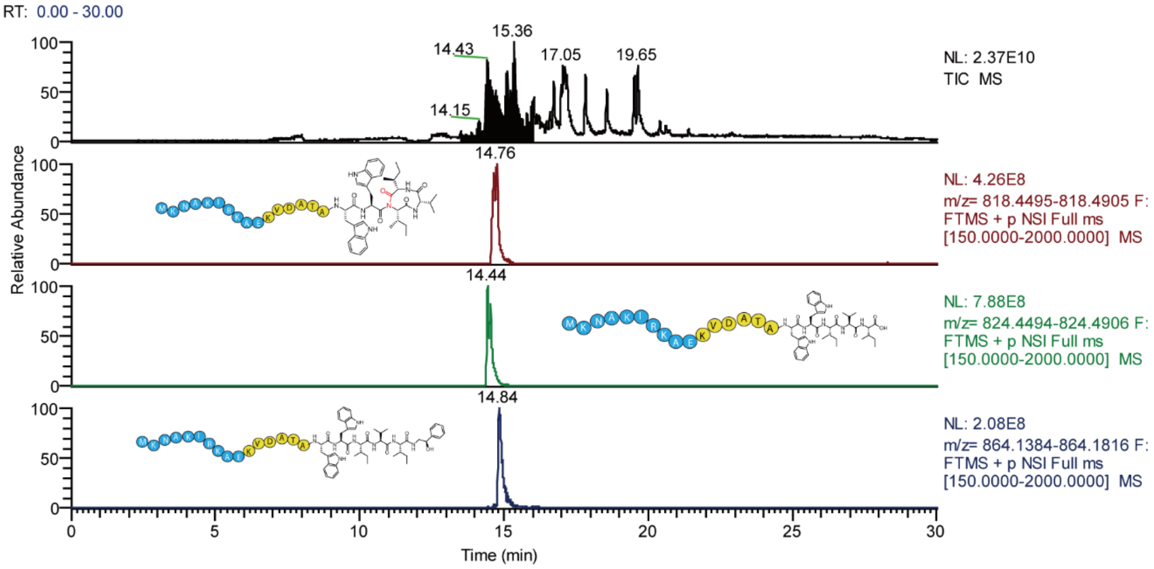


B

### Mass/Charge Table

| | Mass | |
|---|---|---|
| | Mono | Avg |
| (M) | 2589. 45829 | 2591. 05973 |
| (M+H)+ | 2590. 46557 | 2592. 06700 |
| (M+2H)2+ | 1295. 73645 | 1296. 53716 |
| (M+3H)3+ | 864. 16007 | 864. 69388 |
| (M+4H)4+ | 648. 37188 | 648. 77224 |

* C-terminus modification: 119.07

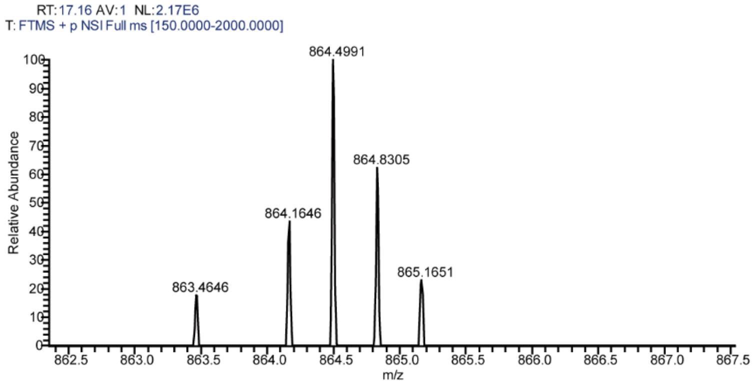


C

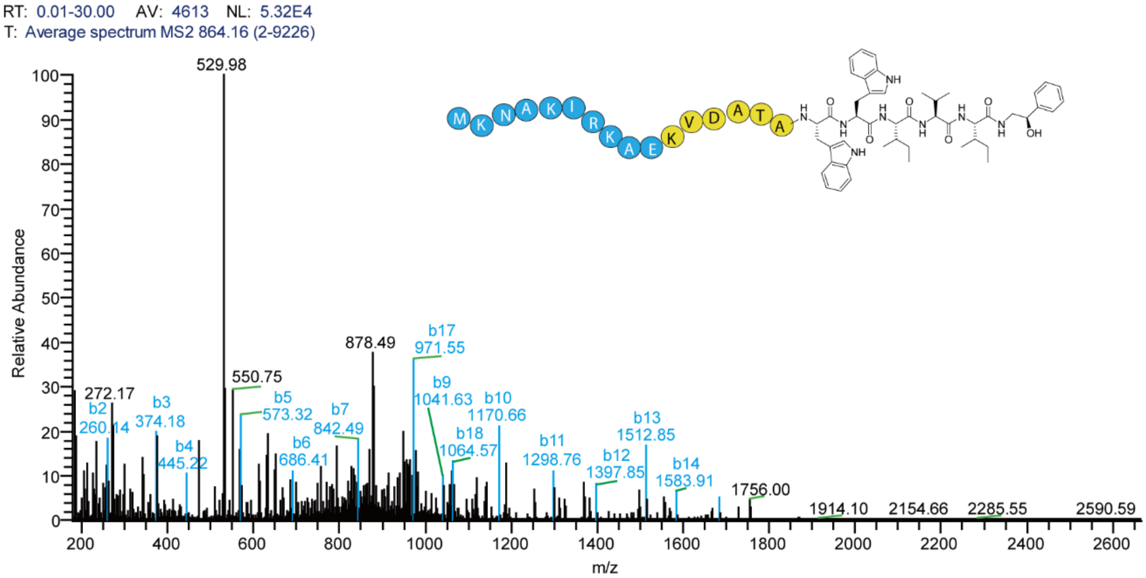


**Supplementary Figure S27.** MS/MS fragmentation of peptides in the enzymatic assays with 2-amino-1-phenylethanol. Fragments of the b-series are highlighted in blue, with those of the y-series in red.

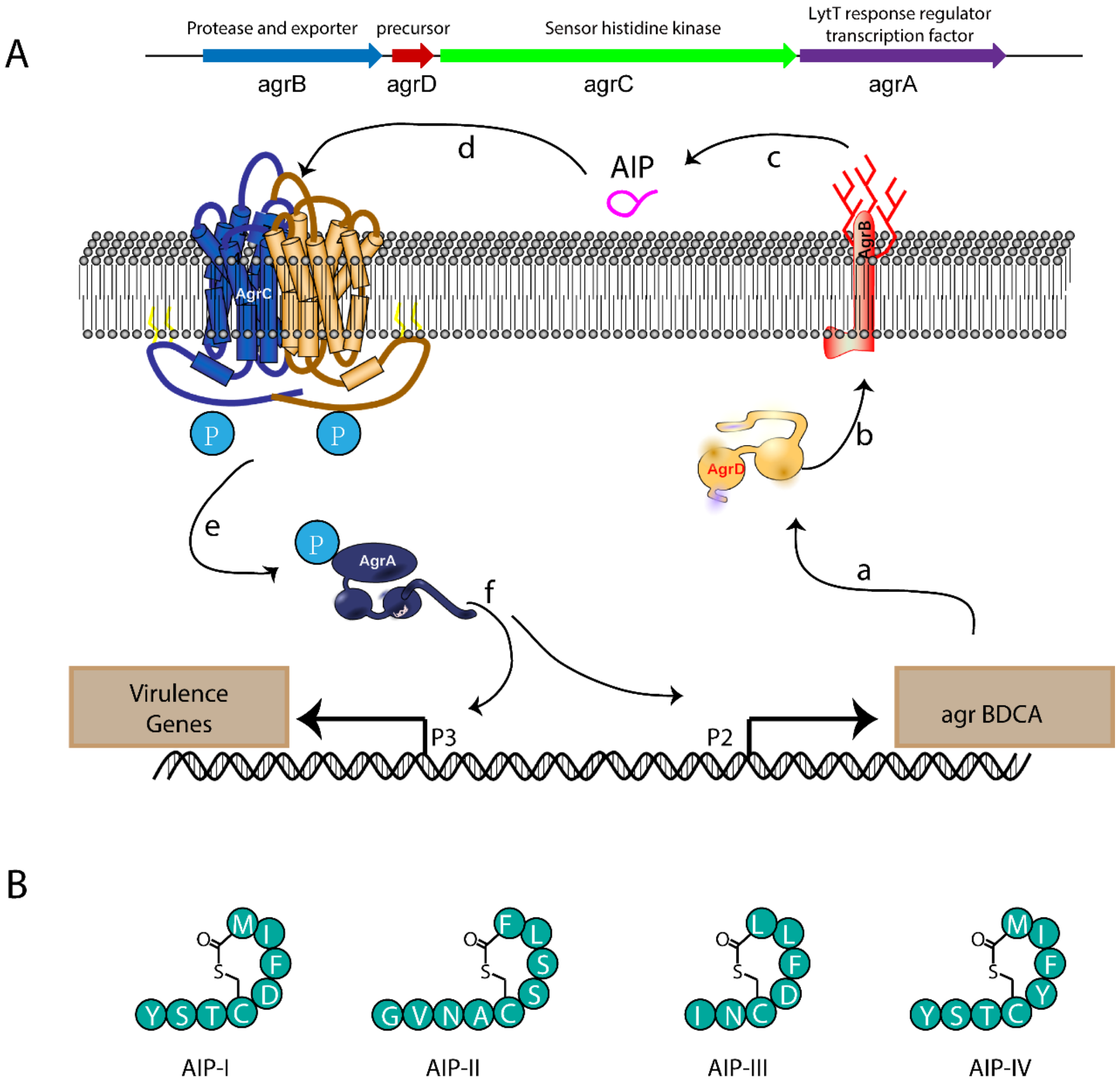


**Supplementary Figure S28.** Schematic of the agr quorum sensing circuit in *S. aureus*. (A) (a) and (b) The precursor peptide AgrD is processed by AgrB. (c) The mature AIP signal is secreted across the cell membrane. (d) AIP binds the extracellular domain of AgrC. (e) The histidine kinase domain of AgrC phosphorylates AgrD. (f) AgrA binds the P2 and P3 promoters to autoactivate the agr system and upregulate RNAIII transcription. (B) Structures and sequences of the four AIP signals (I−IV) corresponding to the four *S. aureus* groups (I−IV). Letters represent amino acid codes.

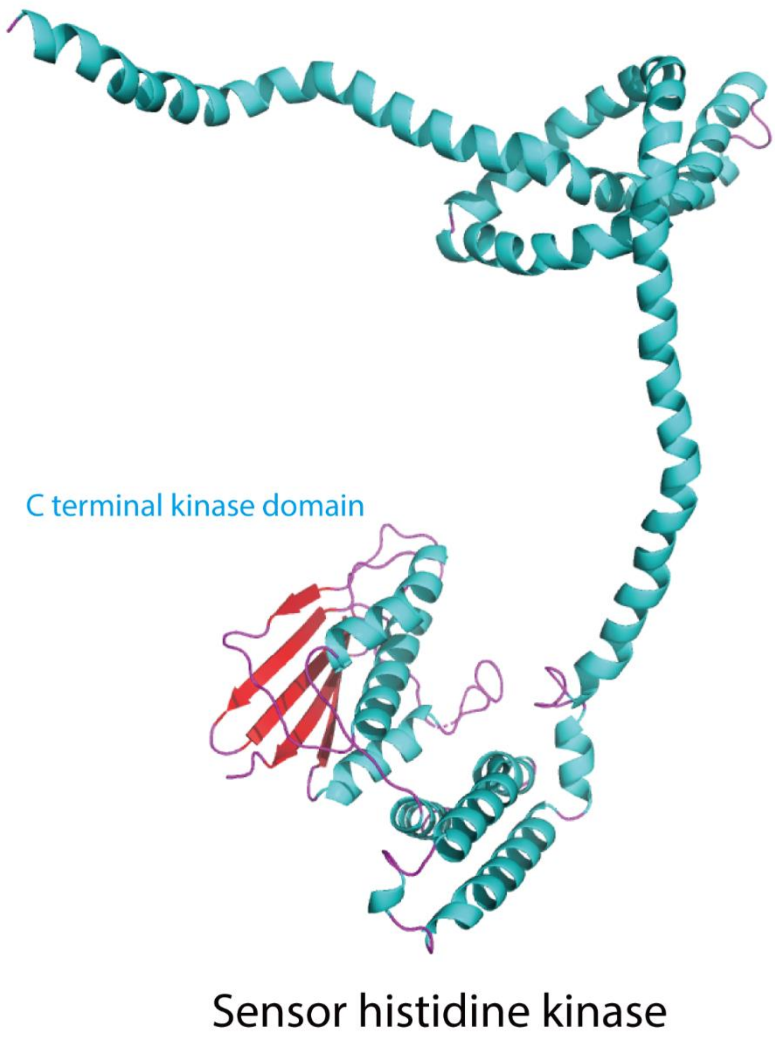


**Supplementary Figure S29.** Homology model of the sensor histidine kinase in the enterofaecin biosynthetic pathway

A

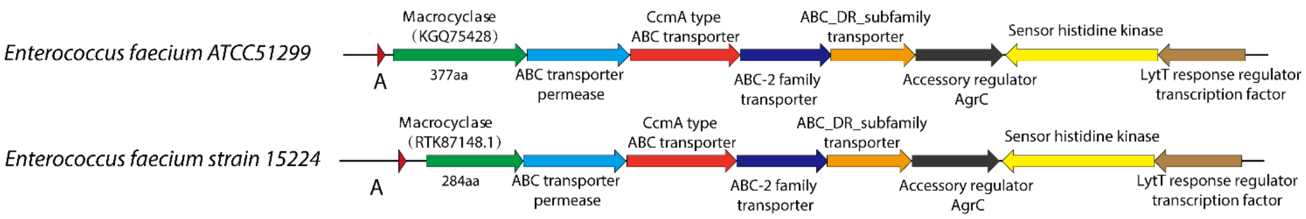


B

| | |
|---|---|
| Program | Blast 2 sequences Citation |
| Query ID | KGQ75428.1 (amino acid) |
| Query Descr | heme biosynthesis protein HemY [Enterococcus faecalis] |
| Query Length | 377 |
| Subject ID | RTK87148.1 (amino acid) |
| Subject Descr | ThiF family adenylyltransferase [Enterococcus faecalis] |
| Subject Length | 284 |
| Other reports | Multiple alignment MSA viewer |

**ThiF family adenylyltransferase [Enterococcus faecalis]**

Sequence ID: RTK87148.1 Length: 284 Number of Matches: 1

Range 1: 1 to 284 GenPept Graphics ▼ Next Match ▲ Previous Match

| Score | Expect | Method | Identities | Positives | Gaps |
|---|---|---|---|---|---|
| 584 bits(1505) | 0.0 | Compositional matrix adjust. | 283/284(99%) | 284/284(100%) | 0/284(0%) |

```
Query  94   MIEDRYLANVNYFSRYCKADDDRFEIQEKINNLKILLLGLGGGGSNILTLLAGLGPKMIR  153
            MIEDRYLANVNYFSRYCKADDDRFEIQEKINNLKILLLGLGGGGSNILTLLAGLGPKMIR
Sbjct  1    MIEDRYLANVNYFSRYCKADDDRFEIQEKINNLKILLLGLGGGGSNILTLLAGLGPKMIR  60

Query  154  MVDYDRVEASNLGRQLLYREADIGEKKTVVAKRAINEMNSNINVETVDKKIIDVNDVVEL  213
            MVDYDRVEASNLGRQLLYREADIGEKKTVVAKR+INEMNSNINVETVDKKIIDVNDVVEL
Sbjct  61   MVDYDRVEASNLGRQLLYREADIGEKKTVVAKRSINEMNSNINVETVDKKIIDVNDVVEL  120

Query  214  TEGIDIIVCAIDEPPFLIHRIVNEAIVKVGLPCVFGASQVSRGRVYTVIPQKTGCFDCMN  273
            TEGIDIIVCAIDEPPFLIHRIVNEAIVKVGLPCVFGASQVSRGRVYTVIPQKTGCFDCMN
Sbjct  121  TEGIDIIVCAIDEPPFLIHRIVNEAIVKVGLPCVFGASQVSRGRVYTVIPQKTGCFDCMN  180

Query  274  LNFSKNDPKFVEQFVGFRNIQFAPPSIAYGPGIFQLTASIVDELIRVVTRYAEPKSLGTQ  333
            LNFSKNDPKFVEQFVGFRNIQFAPPSIAYGPGIFQLTASIVDELIRVVTRYAEPKSLGTQ
Sbjct  181  LNFSKNDPKFVEQFVGFRNIQFAPPSIAYGPGIFQLTASIVDELIRVVTRYAEPKSLGTQ  240

Query  334  YEINYEDGNSFTHKTWPRFESECPTCGKGDVSQWEIFQYYQEKK  377
            YEINYEDGNSFTHKTWPRFESECPTCGKGDVSQWEIFQYYQEKK
Sbjct  241  YEINYEDGNSFTHKTWPRFESECPTCGKGDVSQWEIFQYYQEKK  284
```

**Supplementary Figure S30.** Comparison of the enterofaecin biosynthetic pathway in *E. faecium* strain ATCC 51299 and *E. faecium* strain 15244. A, organization of the gene clusters; B, sequence alignment of the EnfB enzyme in these two gene clusters.

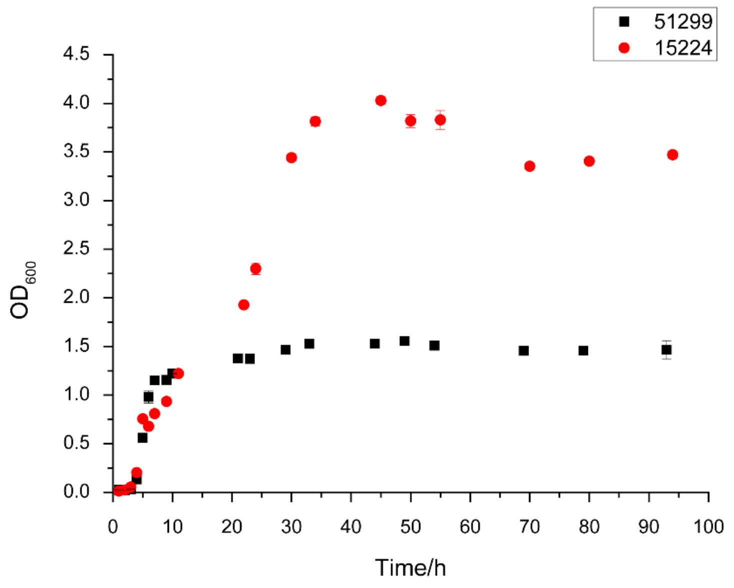


**Supplementary Figure S31.** Growth curve of *E. faecium* strain ATCC 51299 and *E. faecium* strain 15244
